# Imputation integrates single-cell and spatial gene expression data to resolve transcriptional networks in barley shoot meristem development

**DOI:** 10.1101/2025.05.09.653223

**Authors:** Edgar Demesa-Arevalo, Hannah Dörpholz, Isaia Vardanega, Jan Eric Maika, Itzel Pineda-Valentino, Stella Eggels, Tobias Lautwein, Karl Köhrer, Thorsten Schnurbusch, Maria von Korff, Björn Usadel, Rüdiger Simon

## Abstract

Grass inflorescences are composite structures, featuring complex sets of meristems as stem cell niches that are initiated in a repetitive manner. Meristems differ in identity and longevity, generate branches or split to form flower meristems that finally produce seeds. Within meristems, distinct cell types are determined by positional information and the regional activity of gene regulatory networks. Understanding these local microenvironments requires precise spatio-temporal information on gene expression profiles, which current technology cannot achieve.

Here we investigate transcriptional changes during barley development, from the specification of meristem and organ founder cells to the initiation of distinct floral organs, based on an imputation approach integrating deep single-cell RNA sequencing with spatial gene expression data. The expression profiles of more than 40.000 genes can be analysed at cellular resolution in multiple barley tissues using the web-based graphical interface *BARVISTA*, which enables precise virtual microdissection to analyse any sub-ensemble of cells. Our study pinpoints previously inaccessible key transcriptional events in founder cells during primordia initiation and specification, characterises complex branching mutant phenotypes by barcoding gene expression profiles, and defines spatio-temporal trajectories during flower development. We thus uncover the genetic basis of complex developmental processes, providing novel opportunities for precisely targeted manipulation of barley traits.

## Introduction

Plant development depends on the activities of stem cell systems, the meristems, that give rise to organ primordia or new meristems in distinct patterns. In the grass family, flower development involves the sequential specification of meristematic identities at the inflorescence meristem, leading to diverse architectures ^1,2^. Rice and oat plants develop multiple branched panicles, whereas teosinte, rye, wheat and barley form simpler or even unbranched spikes that generate rows of floret meristems, probably reflecting an ancestral state ^3^. There is a strong genetic basis for these inflorescence architectures which can be explored by studying mutants or genetic variation affecting meristem behaviour during domestication. For example, the evolution of maize from teosinte involved the repression of axillary meristems (AMs), enlargement of the inflorescence meristem (IM), and a switch from distichous to spiral inflorescence phyllotaxis, forming multiple kernels at each node ^4^.

In barley, a spike-type inflorescence axis (rachis) generates an AM at each node in a distichous pattern subtended by a transitory and developmentally repressed leaf meristem. AMs differentiate into determinate triple spikelet meristems (TSMs) that separate to form three distinct spikelet meristems (SMs), two lateral (LSM) and one central (CSM) ^5^. Each spikelet initiates a determinate meristem that forms a short axis (rachilla), subtending bracts and a single floret meristem (FM) that generates the floral organs. The fate and determinacy of these different meristems is controlled by genetic and environmental factors. When the indeterminate IM at the tip of the rachis has generated a finite number of TSMs, the IM and several SMs undergo gradual degeneration in a basipetal sequence, indicating that the position of SMs along the rachis is a key determinant of fate. Furthermore, each CSM forms a fertile flower and a single grain, whereas LSMs develop to fertility only in six-rowed barley varieties, but remain small and sterile in two-rowed barley. Mutants have been identified in barley that modulate spike architecture by controlling the fate or identity of specific meristems. For example, LSM fertility and the determinacy of TSMs is regulated by the LOB-domain transcription factor (TF) *VRS4* (*Hordeum vulgare RAMOSA2*, or *HvRA2*) and the TCP-family TF *INT-C* ^6,7^. Loss-of-function mutations in the *INT-M* gene, encoding an AP2-like TF, or in *COMPOSITUM1 (COM1)*, encoding a TB1/CYC/PCF (TCP)-like TF whose expression at the IM-SM boundary depends on *HvRA2*, cause the rachilla to give rise to a new branch or generate more florets per spikelet ^8,9^, indicating that *COM1* promotes meristem determinacy. Mutations in *HvCLV1*, encoding a CLAVATA1 family receptor kinase, or in *HvFCP1*, encoding a secreted signalling peptide that acts through *HvCLV1*, enhance rachilla indeterminacy and fail to maintain *COM2* expression. *HvFCP1* and *HvCLV1* are also required for extended AM initiation at the IM ^10^. At higher temperatures, rachilla determinacy is promoted by the MADS-box transcription factor (MADS-TF) HvMADS33/HvMADS1, a member of the *LOFSEP* clade also associated with inflorescence architectural regulation in rice and several dicot species ^11^.

The homeodomain TF *KNOTTED1* (*KN1*) is a key promoter of meristem indeterminacy during vegetative and reproductive development as first described in maize, and is expressed in the vegetative shoot apical meristem (vSAM) and IM, but not in organ founder cells or leaf primordia ^12^. Only lateral meristems regain *KN1* expression during later development. *KN1* RNA is mostly absent from the outermost cell layer (the L1) of meristems, but KN1 protein together with *KN1*-mRNA can enter the L1 *via* plasmodesmata ^13–15^. Local auxin accumulation combined with the absence of KN1 expression are early markers of organ initiation in meristems of many species ^16^. However, the gene regulatory networks (GRNs) responsible for primordia specification, their identities, determinacy and fates are still largely unknown ^1,2,8,17^. Furthermore, where and when these GRNs execute their function requires precise knowledge on their expression domains.

To explore the GRNs underlying meristem specification and organ initiation in an unbiased manner, we first generated single-cell RNA-sequencing (scRNA-Seq) data for the barley vSAM and inflorescence (spike), thus covering a range of meristem identities and developmental stages. Several scRNA-Seq datasets are available in grass species, providing in-depth expression dynamics for thousands of genes in each cell ^18–25^. Furthermore, deduced developmental trajectories can illustrate shifts in cell fate and concomitant changes in gene expression patterns. However, the origin and fate of cells within complex tissues can only be inferred indirectly from the known expression profiles of prominent marker genes ^26^. Therefore, we precisely localised and quantified the transcripts of 86 genes on tissue sections at cellular resolution using the Molecular Cartography^TM^ (MC) platform for multiplexed single-molecule RNA fluorescence in situ hybridisation (smRNA-FISH). We then integrated the scRNA-Seq and smRNA-FISH data to generate imputed transcriptome-wide single-cell expression matrices. We were able to predict cellular expression patterns with high confidence for most genes and cells on each tissue section. The combined data sets are presented in the user-friendly, searchable online database *BARVISTA* (**bar**ley **v**irtual **i**n **s**itu **t**ranscriptome **a**tlas), which visualises the expression of 48,904 barley genes at cellular resolution on tissue sections representing different developmental stages. *BARVISTA* enables the virtual microdissection of cell-populations from these sections, followed by reclustering and mining for regionally specific gene expression profiles. We used *BARVISTA* to explore developmental trajectories during meristem identity specification and to identify key TFs controlling primordia initiation, meristem formation, identity and determinacy. We show that *HvKN1* transcripts, which are absent from most meristem L1 cells ^14^, can be detected in a subpopulation of L1-cells at the tip of incipient primordia, where *HvKN1* is coexpressed with cytokinin biosynthesis genes, which could establish a positive feedback loop that promotes meristem indeterminacy. We also observed rapid changes in gene expression complexity during SM development and maturation, revealing expression fingerprints that allowed us to phenotype the complex cell identities of meristem mutants. Our integrated dataset, combined with gene expression imputation for vegetative and generative meristem states, revealed gene expression dynamics with unprecedented spatiotemporal resolution, providing a framework for the formation and specification of meristems and organ primordia during barley development.

## RESULTS

### A comprehensive single-cell gene expression atlas of barley meristems

Development of the barley SAM and spike can be described using the Waddington scale (Waddington stages, W) ^27^. From W0 to W1, the vSAM forms lateral leaf primordia in a distichous pattern, which ensheathe the base of the vSAM (**Fig1A,B)**. The vSAM then elongates and transitions to the reproductive stage (IM), characterised by the formation of SMs instead of leaf primordia. By W3.5, the spike has generated two rows of spikelets with a complex developmental gradient. TSMs form from small protrusions on the IM lateral surface (**Fig1C-E**). At P7, the TSM has already separated into the CSM and two LSMs. Further towards the base, P11-P15 are floral meristems (FM) subtended by lemma primordia. P17-P31 have each initiated one carpel and three stamen primordia, but the further formation and differentiation of floral organs is delayed (**Fig1F,G,H,I**). To identify transcriptional changes underpinning the formation of IM, TSM, CSM, LSM, and FM, as well as their developmental stages and the differentiation of organ primordia, we first generated scRNA-Seq data from vSAMs at W0.5 and spikes at W3.5 for the barley reference cultivar Golden Promise (GP). Stage W3.5 captures gene expression information for all meristem types that finally contribute to flower organs (**Fig2A**). Enzymatic digestion of meristem cell walls and the release of viable protoplasts from meristems can be hindered by differences in cell wall composition compared to leaf cells ^28^. We assessed the ability to capture the full diversity of spike cells and tissues by representation of cell/protoplast sizes, comparing intact with protoplasted tissues. We also used the reporter line *pHvFCP1:mVenus-H2B*, which expresses Venus in specific meristematic tissues ^10^ (**SupplFig1A,B,H**). The viability of isolated protoplasts was monitored by differential staining with calcein-acetoxymethyl ester, which fluoresces in the cytoplasm of viable cells following uptake and esterase cleavage, and DRAQ7, which stains nuclei only in membrane-compromised or dead cells ^29^. Only samples with more than 65% viable protoplasts were loaded for microwell-based cell isolation. Cells from inflorescence spikes and vSAMs were isolated in three independent experiments, and we obtained high-quality transcriptome information from 16,528 individual cells for W3.5 ^30^. We detected the expression of an average of 4,527 genes per cell and 17,553 transcripts, mapping to 46,495 gene models, including low and high confidence models, of the reference genome assembly of barley cultivar Morex (MorexV3). Additional libraries from wild-type (WT) vSAM and *com1a;com2g* mutant spikes recovered data from 919 and 5,196 cells, respectively (**Fig2B,SupplTab1**). We then tested how the enzymatic protoplasting protocol affected gene expression by comparing bulk RNA-Seq data from intact inflorescence tissue and protoplasted tissues, using three biological replicates. We identified 3,684 genes that were differentially expressed in response to protoplasting using a log fold change (log_2_FC) ≥ 3 for upregulated genes and ≤ −3 for downregulated genes and a false discovery rate (FDR) < 1×10^−5^, (**SupplFig1C-G,SupplTab2**). We then subtracted the protoplasting-induced differentially expressed genes (DEGs), and integrated the scRNA-Seq data for the three independent experiments using the most variable 7,500 genes, followed by dimensional reduction for clustering analysis. We identified 27 clusters with 4,256 putative unique markers genes (**Fig2A-D,SupplTab3**). Cluster ontogenies were determined using known marker genes (**SupplTab4**).

**Fig1:**
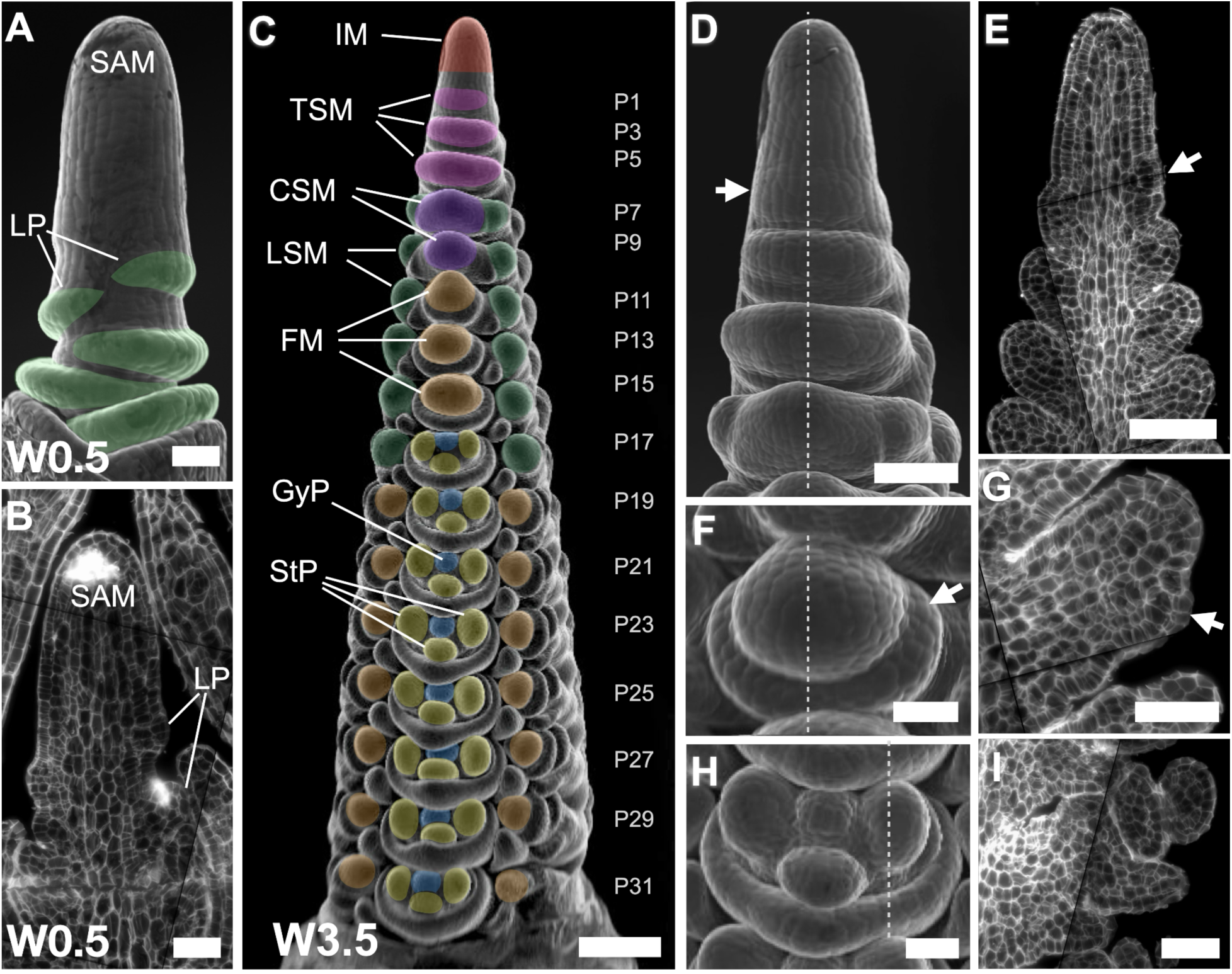
Organ formation in developing spikes of barley. (A) Scanning electron micrograph (SEM) of the vegetative shoot apical meristem (vSAM), highlighting leaf primordium (LP) formation (green). (B) Sagittal section of the vSAM. (C) At W3.5, the inflorescence meristem (IM) at the top of the spike is determined to form lateral meristems specified as triple spikelet meristems (TSMs), which will then generate two lateral spikelet meristems (LSMs) and one central spikelet meristem (CSM). In the Golden Promise genetic background, a two-row type of barley, only the CSM continues its development and will form a floret meristem (FM), which then generates gynoecium (GyP) and stamen primordia (StP). Plastochrons are indicated as P1–P31. (D) SEM of the upper part of a developing spike, where the TSM is specified and apparent as a protrusion on the surface axillary to the IM (arrow). (E) A sagittal section through the IM and TSM. The earliest sign of the TSM is cell proliferation where the primordium is formed (arrow). (F) SEM of a FM with lemma primordium (arrow). (G) FM sagittal section showing the outgrowing lemma (arrow). (H) SEM of one floret developing StP and GyP. (I) Sagittal section through one floret. Dotted lines in D, F and H are equivalent in position to the sagittal sections in E, G and I, respectively, Scale bars: (C) 200 µm; (D) and (E) 100 µm; (A), (B) and (F)-(I) 50 µm.

**Fig 2:**
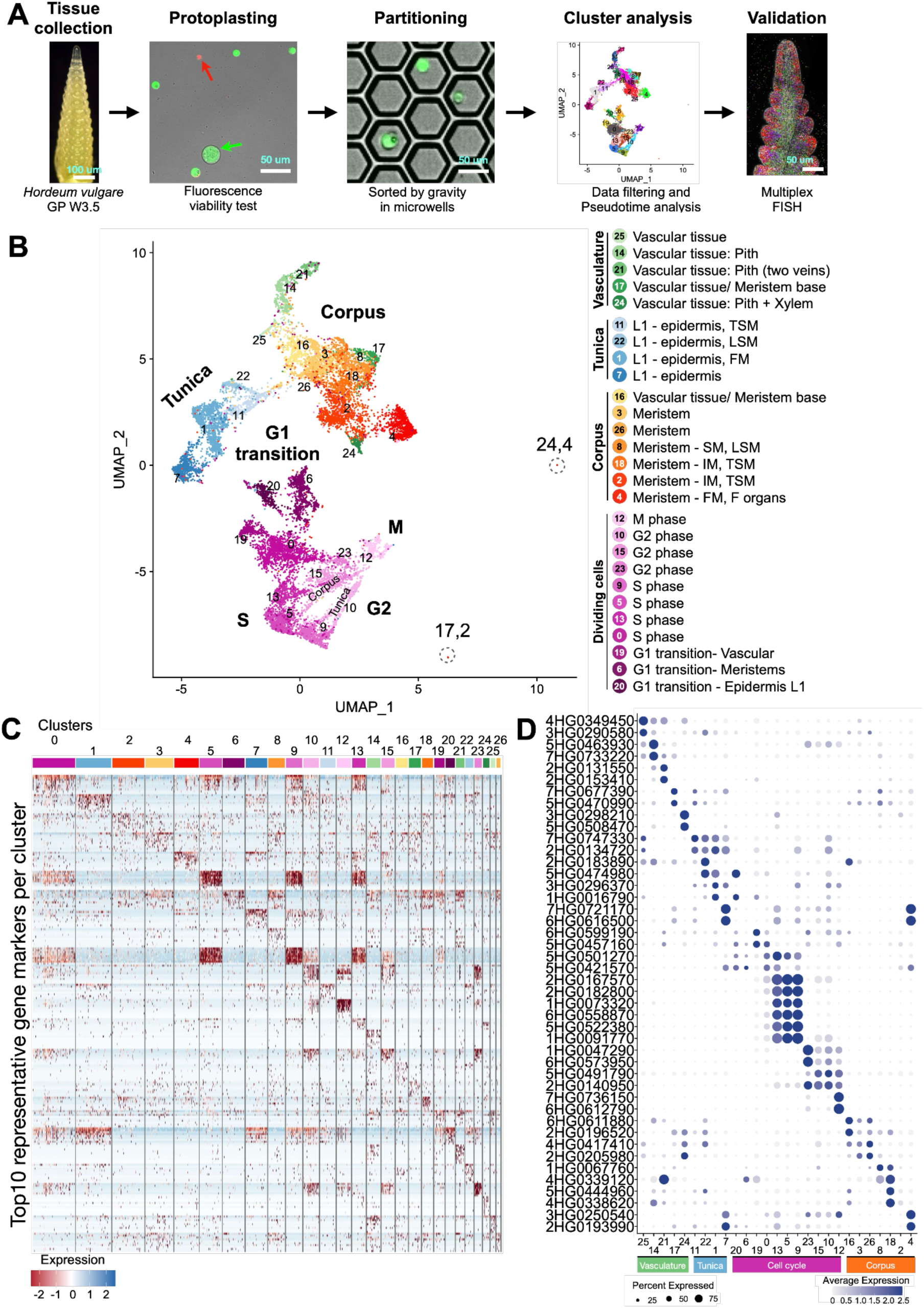
A single-cell atlas from developing barley spikes. (A) Workflow for the isolation, sequencing, analysis and validation of protoplast scRNA-Seq from developing barley spikes at WS3.5 (inflorescence meristem, IM; triple spikelet meristem, TSM; central spikelet meristem, CSM; lateral spikelet meristem, LSM; floret meristem, FM; stamen primordia, StP; gynoecium primordia, GyP). (B) Uniform manifold approximation and projection (UMAP) for dimension reduction revealed that the cluster identities are organised as four populations containing vasculature, tunica, corpus, and dividing cells. (C) Heat map of 10 representative marker genes per cluster. (D) Expression of two representative marker genes organised by cell populations. Scale bars: (A) 200 and 50 µm.

### scRNA-Seq identifies markers specific for tissue lineages

We identified four distinct populations of cells: vasculature, tunica, corpus and dividing cells (**Fig2B-D,SupplFig2**). Each population comprised multiple clusters, clearly separated by groups of cells potentially derived from specific meristem types. We first used *HvKN1* (4HG0339120) to identify clusters of meristematic and vascular tissues. A complementary group of clusters representing the tunica expressed *HvHDZIV8* (2HG0198150), which encodes a homeodomain-leucine zipper TF. *HDZIV8* is related to the maize *OUTER CELL LAYER* TFs specific to the tunica layers, and the Arabidopsis L1 marker *MERISTEM LAYER1* (AtML1) ^31^. Additional marker genes identified specific subpopulations in the tunica and corpus: for the IM and TSM, *HvMADS42*/*MADS34* (5HG0511250); for the lemma and CSM, *HvMADS33*/*MADS1* (4HG0396400); for the CSM and floret organs, *HvMADS68*/*MADS7* (7HG0684020); for TSM founder cells, glumes and LSM, VRS4 (3HG0233930); for the floret meristem, *HvMADS50*/*MADS6* (6HG0604360); and for stamen primordia, *HvMADS24*/*MADS2* (3HG0307160) and *HvMADS76*/*MADS16* (7HG0721170). Similarly, markers were identified for the vascular tissues: cambium, *HvPXY* (7HG0701280); xylem, *HvVND1* (4HG0410880); phloem, *HvSMXL3* (2HG0167230); and rachis, *HvPSAN* (2HG0113750). For the cell-cycle, we identified S-Phase histone markers, *H3* (7HG0658120), *H4* (6HG0550690), *H2A* (5HG0518610), and *H2B* (1HG0067590); G2-phase markers *Cyclin-A* (3HG0249410) and *CDKB* (7HG0680610); and M-phase markers *HvTANGLED1* (1HG0073890), closely related to maize *TANGLED1* encoding a microtubule-binding protein associated with mitotic division ^32^; *EARLY NODULIN-LIKE PROTEIN* (1HG0082450) and eisosome protein (4HG0411400) (**SupplFig2, SupplTab4**). Our diverse dataset ranged from cell-lineage initials to differentiated cells. Based on 7,500 variable genes, we captured cell-cycle specific transcriptomes for the tunica and corpus, as well as phase-specific markers. To avoid biases due to cell-cycle phases, we then focused on cells in G1.

### Combination of smRNA-FISH and scRNA-Seq identifies meristem subdomains

We validated our cluster annotation of meristem domains by smRNA-FISH (**SupplTab5**, **SupplFig3**) ^33^. To simplify MADS-box nomenclature, we use the most recent barley (HvMADS) ^34^, but genes previously named by homology to characterised rice genes are shown with both names (MADS, also listed in **SupplTab6**). Marker gene expression profiles in the developing spike can define the sequence of gene activation during the specification of meristem-types, enabling the identification of novel gene expression domains preceding early differentiation. We selected 100 genes from the scRNA-Seq data, (3-4 per cluster) and generated multiplex smRNA-FISH probes for hybridisation to tissue sections representing different barley developmental. For W3.5 we reliably detected transcripts from 86 genes, with a spatial resolution of <140nm. For example, *HvHDZIV8,* two *HDZIV*-related TFs (7HG0702200 and 6HG0618540) and a pleiotropic drug resistance ABC transporter gene (3HG0240110) were specifically expressed in the tunica of the vSAM (**Fig3A, SupplTab7**) and IM at stage W3.5 (**Fig3B**), whereas *HvKN1* transcripts were mainly expressed in corpus cells. To determine whether the smRNA-FISH data reflected overall expression levels, we compared the total number of RNA-Seq reads from W3.5 spikes with the number of spots detected by smRNA-FISH. Over a range of genes with diverse expression levels, we observed a Pearson correlation of 0.73 between the two datasets, indicating that the number of smRNA-FISH dots is a reliable proxy for gene expression levels (**SupplFig4A**). Following cell segmentation, we assigned counts per gene and per cell and generated an expression matrix for the 86 genes detected across 22,082 segmented cells, with a dynamic range of up to 536 transcripts of a single gene per cell, a maximum of 627 total transcripts per cell and a mean of 95.35 transcripts per cell (**SupplFig4B-E, SupplTab5**).

**Fig3:**
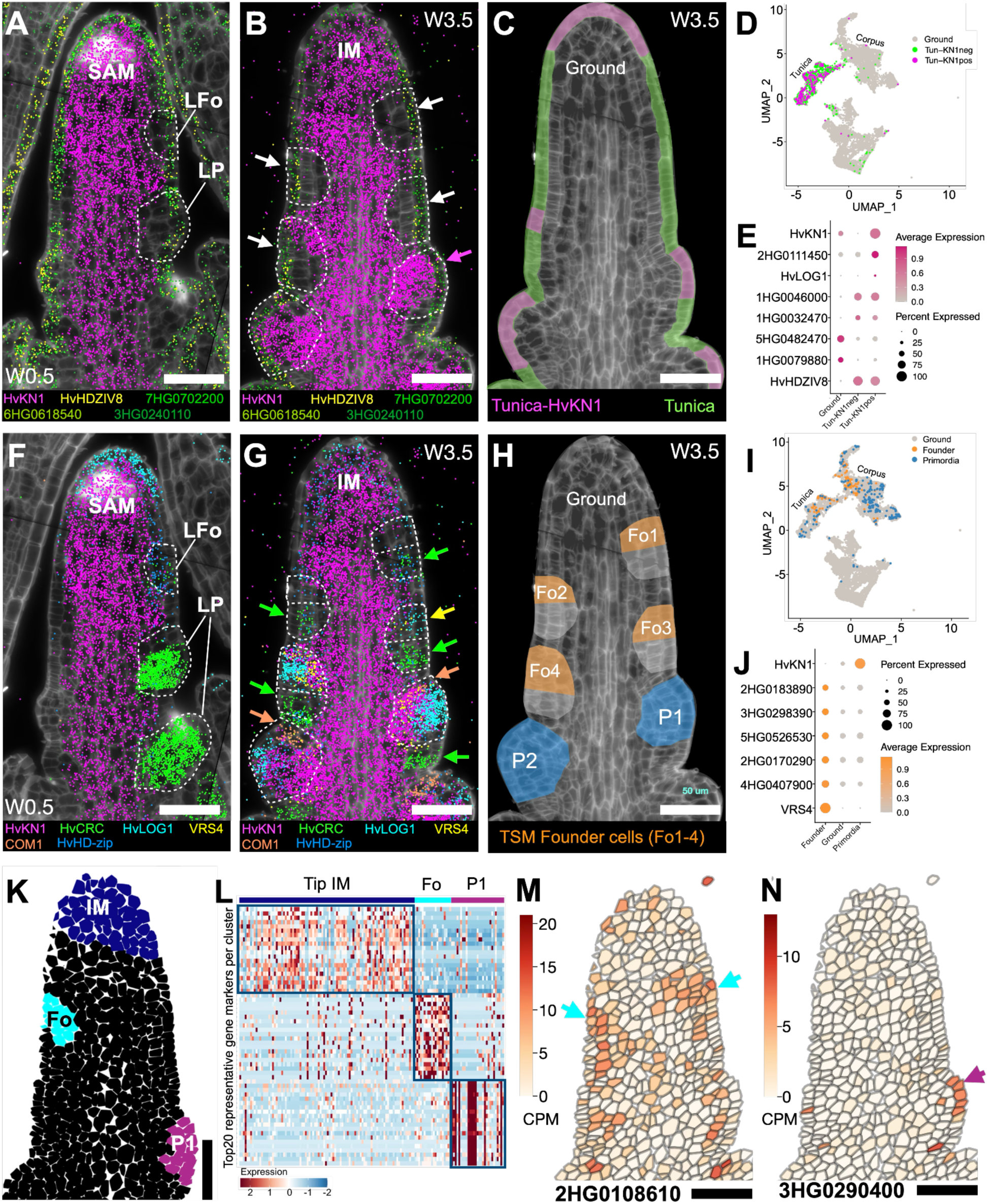
Identification of genetic determinants in primordia and early meristem specification. (A) In vegetative SAM (vSAM) the formation of leaf primordia (LP) is preceded by the depletion of *HvKN1* mRNA (magenta dots) in the leaf founder cells (LFo) and during LP growth. (B) In inflorescence meristems (IM), spikelet founder cells (white arrows, and interrupted line) also show *HvKN1* mRNA depletion, but this is restored once the primordia are organised (magenta arrow). (C) The organisation of tunica and corpus in the IM is reflected by their expression profiles, but in developing spikes we see an overlap of *HvKN1* in the tunica, especially in meristem tips. (D) Differentially expressed genes in tunica cells expressing *HvKN1*. (E) Candidate marker genes in tunica cells expressing *HvKN1*. (F) In the SAM, *HvHD-zip* and *HvCRC* are expressed in the leaf primordia founder cells (LFo), but in the established leaf primordia (LP) only *HvCRC* remains strongly expressed with no restoration of *HvKN1* in the primordia. (G) In spikelet founder primordia, where *HvKN1* mRNA is depleted, the suppressed bract is pre-patterned (green arrow) and expressing leaf-like marker genes such as *HvCRC*, and the spikelet meristem founders cells (Fo) can be identified by *VRS4* and *HvLOG1* expression (yellow arrow). One of the earliest signs of triple spikelet meristem primordia is the expression of *HvCOM1* (orange arrows). (H) Schematic representation showing the spikelet meristem founder cells (Fo) and the triple spikelet meristem primordia (P) in a developing spike of barley. (I) Founder (Fo) and primordial (P) cells can be subtracted from scRNA-Seq populations using their expression profiles. (J) Candidate genes differentially expressed in triple spikelet meristem founder cells. (K) By data integration and imputed gene expression, we can virtually dissect the IM (dark blue), Fo (cyan) and P cells (purple). (L) Heat map of top 20 genes expressed differently in each group of cells. Heat maps show selected examples of imputed differentially expressed genes using a Model-based Analysis of Single-cell Transcriptomics (MAST), with *p_adj_* < 0.05 and with an enrichment of cells expressing the genes in the analysed population more than double the level in the remaining cells (pct.1/pct.2 ≥ 2). (M) Imputed expression of candidate genes specifically expressed in the Founder cells or (N) in TSM primordium. Scale bars = 50 µm.

### Integrating scRNA-SEQ with smRNA-FISH data for gene expression imputation

The position of a cell within a tissue and its gene expression profile may provide information on developmental time and progression, but also the clonal origin of a cell. Current technologies for quantitative spatial gene expression analysis are limited by the number of probes, sensitivity in molecularly crowded environments, and the retention or transferability of RNA molecules to capture devices ^33,35^. We therefore used the spatially resolved quantitative data as anchors and mapped each cell in the scRNA-Seq dataset to its nearest neighbour in the MC dataset. Of the 86 genes in the MC dataset, 81 were expressed in a sufficient number of cells and at differing levels to be regarded as informative. Integration of the datasets in *Seurat* ^36^ following the normalisation of gene expression values yielded clusters, whose numbers depended on the *clustering resolution* (*cr*) parameter, which we tested at settings of 0.1 to 2.0. Biologically relevant clusters representing “dividing cells”, “floret meristem”, “xylem” and “L1 – epidermis” were identified at *cr0.3,* indicating that these involve highly distinct gene expression patterns, separating them from other cell populations (**SupplFig5**). “L1 – FM epidermis”, “L1 – F organ epidermis”, and “early TSM” became distinct only beyond *cr1.1*, indicating fewer differences in gene expression relative to other cell populations.

We developed a data imputation approach to map gene expression values from the scRNA-Seq data onto MC cells. In the initial cell similarity determination step, proximate cells from the scRNA-Seq experiment were determined for each MC cell using cosine similarity (CS) ^37^ (**SupplFig6A**). Gene expression values for each MC cell were then calculated as a weighted average from the 25 most similar scRNA-Seq cells, using the CS as weight to ensure that the most similar cells had the greatest influence on the calculated average (**SupplFig6B**). Expression levels were imputed for 48,904 genes in the inflorescence sections and 27,656 genes in the vSAM. To validate the accuracy of the method, we compared imputed and experimentally determined gene expression values (**SupplFig6C**) on a cell-by-cell basis for all 22,082 segmented cells, using expression data for the anchor genes to determine the CS between cells. A CS of +1 indicates that the entire gene expression profile of a cell can be computationally reproduced. Mean CS values were 0.84 ± 0.11 for the inflorescence sections and 0.78 ± 0.11 for the vSAM, showing that gene expression patterns in most cells could be reconstructed with only minor differences compared to measured values (**SupplFig7**). Because the 81 anchor genes were used for both initial cell similarity determination and validation, we performed simulations to validate the quality of imputation. The scRNA-Seq data were split into a test set, for which gene expression was imputed (replacing the MC dataset in the workflow described above), and a reference dataset from which gene expression data were taken. For both imputations, the 81 anchor genes were used during the cell similarity determination step. For the first imputation, validation involved the same 81 genes (scSub IM 81 genes) (**SupplFig8**). For the second validation, all 48,904 genes (scSub IM 48,904 genes) were used for the cell-by-cell comparison of imputed *vs.* experimentally measured values. Validating with the same 81 anchor genes gave a mean CS of 0.84 ± 0.10, and validation with all 48,904 genes gave a still robust, although slightly lower CS of 0.73 ± 0.10 (**SupplTab8**). We then explored the impact of cell number and gene selection on the accuracy of the imputation. Accuracy improved linearly with increasing reference cell number, but plateaued at ∼2,500 cells (**SupplFig9**). We also found that genes with substantial variations in expression across the cell samples performed better than less variably expressed genes. Genes selected using *PERSIST*, which identifies representative gene sets for distinct cell populations ^38^, outperformed hand-selected or randomly selected genes, and 20 genes were sufficient for reliable cell similarity determination.

Results are available through the online database *BARVISTA,* which displays individual cell clusters, uniform manifold approximation and projection (UMAP) plots and imputed gene expression data for all cells in a tissue section. Individual cells or groups can be selected to extract gene expression data for further analysis.

### Exploring gene expression programs during primordia initiation

We used our integrated datasets in *BARVISTA* to probe gene expression patterns in detail during early primordia initiation in the vSAM or IM. *KN1* is often used as a marker for the meristem corpus, which is mostly absent in the L1 layer and downregulated at sites of organ initiation ^14^. However, scRNA-Seq data in maize identified a small number of tunica cells expressing *ZmKN1* ^39^. The rice ortholog of *ZmKN1* (*OsH1)*, is expressed in the L1 of floret meristems, in some cells of the IM, but not in L1 cells of the vSAM ^40^. The function and origin of *KN1* RNA, which might bind to mobile KN1 protein in the corpus for co-transport to the tunica, is unknown. We used the expression profiles of *HvKN1* and *HvHDZIV8* to identify the tunica cells in our scRNA-Seq expression data (**Fig3A,B**). In the spike, almost 9% (268 of 3,023) of all tunica cells expressed *HvKN1* (**Fig3C,D**). Differential expression analysis revealed that the *HvKN1*-positive tunica cells also contained RNA from the cytokinin biosynthesis gene *HvLOG1* (5HG0502720), as well as *HvWOX9C.2* (3HG0278560) and *HvWOX9C.1* (1HG0088440), which are *WUSCHEL-LIKE HOMEOBOX* genes related to rice *OsWOX9C*. Interestingly, *HvLOG1* is expressed at the tips of the SAM, IM, TSM and FM (**Fig3E, SupplTab9**), suggesting that it might generate a cytokinin source at meristem apices and promote *WOX9C* expression. We extended our analysis to the vSAM (**SupplFig10A-D, SupplTab10**). Here, ∼12% (17 of 138) of all tunica cells expressed *HvKN1* (**SupplFig10C,D**). Among genes differentially expressed between the corpus tissue and tunica clusters, we found an ARGONAUTE gene (2HG0125150) related to *OsAGO14* (**SupplFig10D**). However, smRNA-FISH indicated minimal expression of *HvKN1* in tunica cells of the vSAM. *HvKN1* expression patterns in the tunica resemble those of *OSH1* in rice, with minimal expression in the vSAM, but higher levels in reproductive meristem tips.

*HvKN1* is expressed in the IM but is absent in founder cells (Fo) of the TSM primordium, providing an early marker for Fo cell specification (**Fig3B,F-H**). The TF encoded by *HvHD-zip*/*HvHDZIV2*/*HvOCL2* (2HG0187460) was previously reported as an IM or spike tip marker, but our smRNA-FISH data showed it is also expressed at the tips of other meristems within the inflorescence ^41^ (**Fig3F,G**). Prepatterning of Fo cells into TSM and suppressed bract becomes apparent from Fo3 onwards. TSM founder cells are marked by the downregulation of *HvKN1* and concomitant expression of *VRS4* (3HG0233930) and *HvLOG1* (5HG0502720), or by the downregulation of *HvKN1* and upregulation of *CRABS CLAW* (*HvCRC*, 4HG0396510) where the suppressed bract will form (**Fig3B,G,H**). Once the TSM visibly protrudes from the IM, *HvKN1* and *HvHD-zip* (*HvHDZIV2* or *HvOCL2*) are re-expressed within the TSM. Within the growing TSM, the expression of *HvCOM1* (5HG0479720), encoding a TCP-like TF, marks the formation of the rachilla (**Fig3B,G,H**). Using these expression profiles, we were able to isolate the TSM founder cells (Fo) (**Fig3I**) and identify genes specifically expressed in these cells, including an Argonaute gene (4HG0334490), and those encoding HD-ZIP (6HG0611880), MADS-box (*HvMADS84*, 1HG0054230), AT hook motif DNA-binding (7HG0647300) and MYB (7HG0698540) TFs (**Fig3J,SupplTab11**). We used smRNA-FISH data to characterise the TSM Fo in detail (**Fig3G,H,K**), and isolated Fo, IM and P1 cells by virtual microdissection in *BARVISTA*. The analysis of DEGs identified candidates that expressed in each cell population (**Fig3L-N,SupplTab12**), including 27 expressed specifically in Fo cells. The latter included DEGs encoding a leucine-rich repeat protein kinase-like receptor (2HG0108610, **Fig3M**), two BREVIS RADIX-like transcriptional regulators (2HG0193530 and 5HG0535760) ^42^ and two YABBY TFs (*YABBY1*/*TONGARI-BOUSHI1-related*, 2HG0184460 = *HvTOB1*, and *YABBY15*/*TOB2-related*, 6HG0598850 = *HvTOB2*), both involved in node/internode specification in rice, and regulated by the homeodomain TF OsH1 ^40^. *HvTOB2* was detected using both approaches, suggesting YABBYs are involved in the specification of TSM Fo cells. Among the markers for IM, we found *HvFT2* (3HG0244930), a homolog of *OsFT-L1*, which is activated in rice IM by the mobile paralogs *OsHd3a* and *OsRFT1*. *OsFT-L1* is also expressed in the undetermined primary and secondary branch meristems, but not in the mature IM ^43^. The function and expression profile of *HvFT2* in developing barley spikes are unclear. Another flowering regulator, the bZIP TF *HvFD7* (2HG0111000) was enriched in the IM like its rice ortholog *OsbZIP62*, which physically interacts with OsFT-L1 ^44^. Primordia also express one of the two *AtSCR* homologues, *HvSCR1* (4HG0353780). In Arabidopsis, *AtSCR* is involved in the establishment of primordia and is expressed in FMs as early as P1 ^45^.

### Gene expression programs governing meristem and floral organ identity

Subclustering in the vasculature revealed cell populations not previously detected by our general analysis (**Fig2B**), including phloem cells (**SuppFig11**, **SuppFig12**, **SupplTab13**). We therefore analysed meristem-to-organ transitions in developing spikes, focusing on the highly diverse corpus cells, thereby avoiding developmental trajectories in multiple directions (e.g. leading from initial cells to vasculature, tunica or corpus cells simultaneously) (**Fig2B-C**,**SupplFig2,3**). Within the corpus, we identified 18 new subclusters, including those annotated as TSM, CSM, FM and the primordia of developing floret organs, such as glumes, lemma, palea, lodicules, and developing stamens (**Fig4A**, **SupplFig13A**, **SupplTab14**). To identify genes modulated along the developmental trajectory from IM to floret organs, we applied pseudotime analysis starting from cells expressing *COM1* and *VRS4* (subcluster 16, **Fig4A,B**, **SupplFig13B-C**), as bona fide markers for the establishment of SMs (**Fig3G**). The inferred pseudotime trajectory was used to track the establishment of CSM and LSM from TSM, from CSM to FM, and from FM to organ development (**Fig4B**, **SupplFig13D-S**). We then searched for potential master regulators of developmental progression by focusing on key TFs whose expression commences at the branch points of the pseudotime trajectory. Among the genes detected in corpus subclusters, we found 46 MADS TF genes, 26 of which were differentially expressed between the 0.24-74.7% of the 4807 cells in the corpus (**Fig4C-D**, **SupplTab15**). *HvMADS42*/*MADS34* and *HvMADS64*/*MADS5* are activated early and mostly coexpressed in the IM and TSM. *HvMADS42* expression declines following the differentiation of the CSM and later localised to the adaxial side. *HvMADS64* and *HvMADS33* are both expressed in the CSM, but only *HvMADS33* will also mark the developing lemma and palea. *HvMADS50*/*MADS6* expression arises in the established FM and, like *HvMADS33*, continues in the lemma and lodicule primordia. *HvMADS40*/*MADS8* is expressed in floret organ primordia, including lodicule, stamen and gynoecium primordia (**Fig4C-D**,**Fig5A,B**). Twelve MADS TF genes were strongly modulated by pseudotime (Moran’s I ≥ 0.1 and q ≤ 3.4×10^−162^, **SupplFig14**, **SupplTab16**) and their expression profiles correlate with their developmental activation, thus highlighting two axes, one from the apex to the base of the spike, and another from the base to the tip in the FM developing floret organs. According to pseudotime (pT) analysis (**Fig4F**), we identify six trends in gene modulation, with gene expression peaking around major meristem transitions such as IM to TSM (pT0.5), TSM to SM (pT5), SM to FM (pT9), FM to Lemma and Palea primordia (pT14) and two from FM to stamen, gynoecium and lodicule primordia, one group of genes starting earlier than the second group, but both having a late peak (pT18) (**Fig4C-F**). Clustering of modulated genes by pT revealed two main clades: those expressed between IM to FM, and those involved later in floret organ organisation. The first clade showed more complex gene expression patterns (IM to FM) than the second (FM to floret organ formation), whereas organ development involved more modulated genes but only two patterns in late modulation (pT >14 or >18) (**Fig4E).** We found that 20 of 26 MADS TF genes were modulated during and after the establishment of the FM, consistent with the proposed role of these TFs to define the floral whorls after floret transition (**SupplFig14**). Based on the pT activation profiles, we can infer which other MADS TF genes are expressed in specific floral whorls (**Fig5,SupplFig14**). Grasses follow the ABCDE model of floral development, where organ determination is mediated by MADS-TF multimers ^46,47^. By analysing the detailed expression profiles of 12 MADS TF genes, we were able to assign a putative A, B, C or E class to all such genes expressed in corpus subclusters and modulated by pT, taking as a reference the proposed model for rice ^48^ (**Fig5C,D,SupplFig14**). Based on genes modulated during major transitions, we can correlate pT and canonical time expressed as plastochrons (**Fig4F**). Within corpus subclusters, 4733 cells expressed at least one of the 26 HvMADS-TF genes (median = 5). By analysing the presence or absence of these HvMADS-TF genes and their pT classes, we could then assign each cell to specific organs within florets (**SupplFig15**), also revealing the transcriptional profiles for specific floral organs at incipient stages.

**Fig4:**
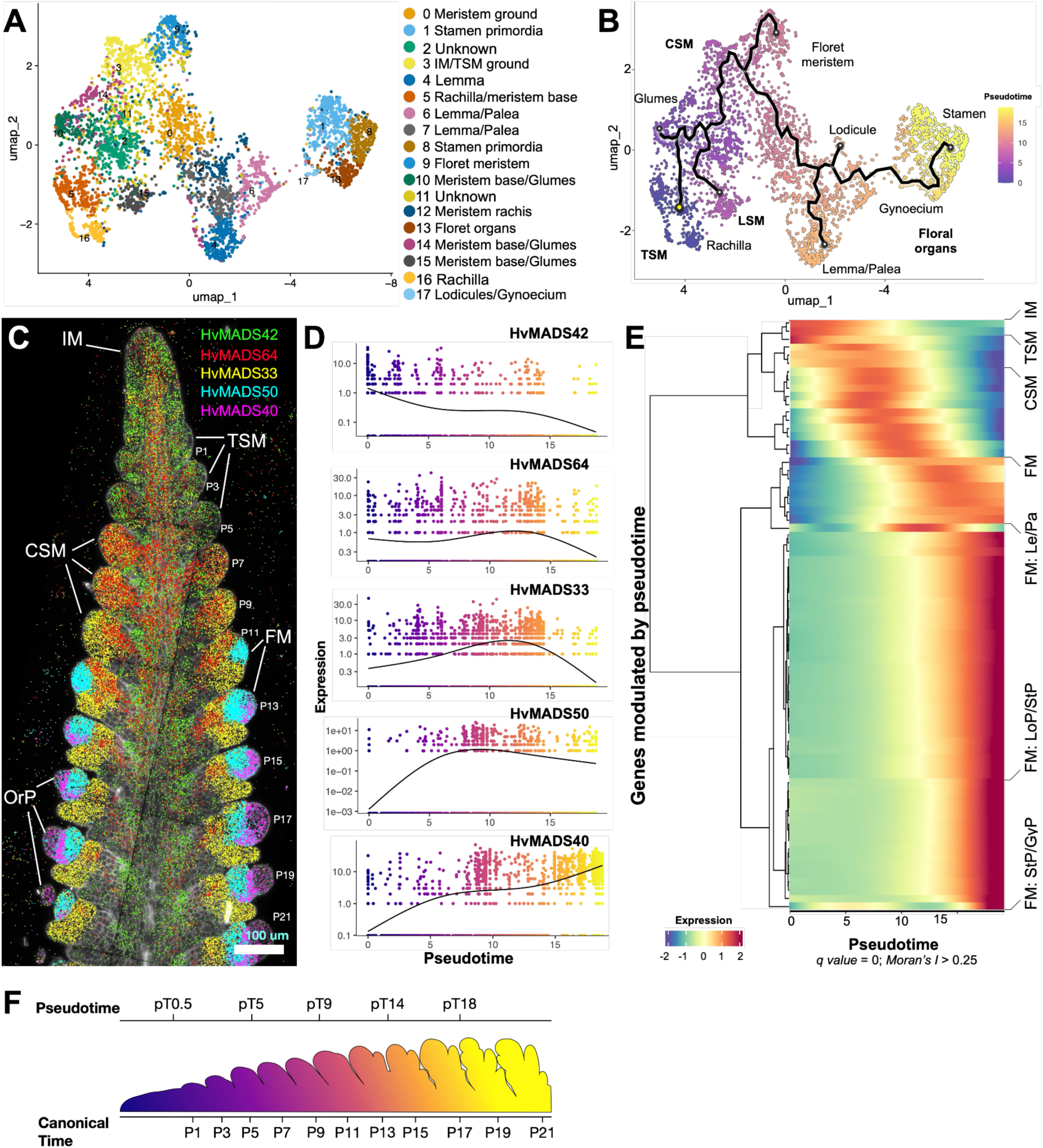
Subclustering in corpus cells postulates MADS-box transcription factor genes as markers for specific meristems and organ formation. (A) UMAP and subclustering of corpus cells from developing spikes in barley. (B) Pseudotime analysis in subclusters from corpus cells defines the formation of meristems and organs from inflorescence meristem to organs in the floret meristem (triple spikelet meristem, TSM; central spikelet meristem, CSM). The yellow dot shows the starting point in pseudotime, the black line shows the developmental trajectory across the pseudotime, and endpoints for primordia formations are shown as circles. (C) Expression profiles of selected MADS-box transcription factor genes along the developing spike (inflorescence meristem, IM; floret meristem, FM; floret organ primordia, OrP). (D) Expression profile according to pseudotime analysis for selected MADS-box transcription factors. (E) Genes with the highest differential expression according to pseudotime analysis grouped by hierarchical clustering (Moran’s I ≥ 0.25 and q = 0), showing the major developmental transitions during spike development. (F) Inferred equivalency of pseudotime and canonical time, as a plastochron, based on genes modulated by pseudotime. Scale bar: (C) 100 µm.

**Fig5:**
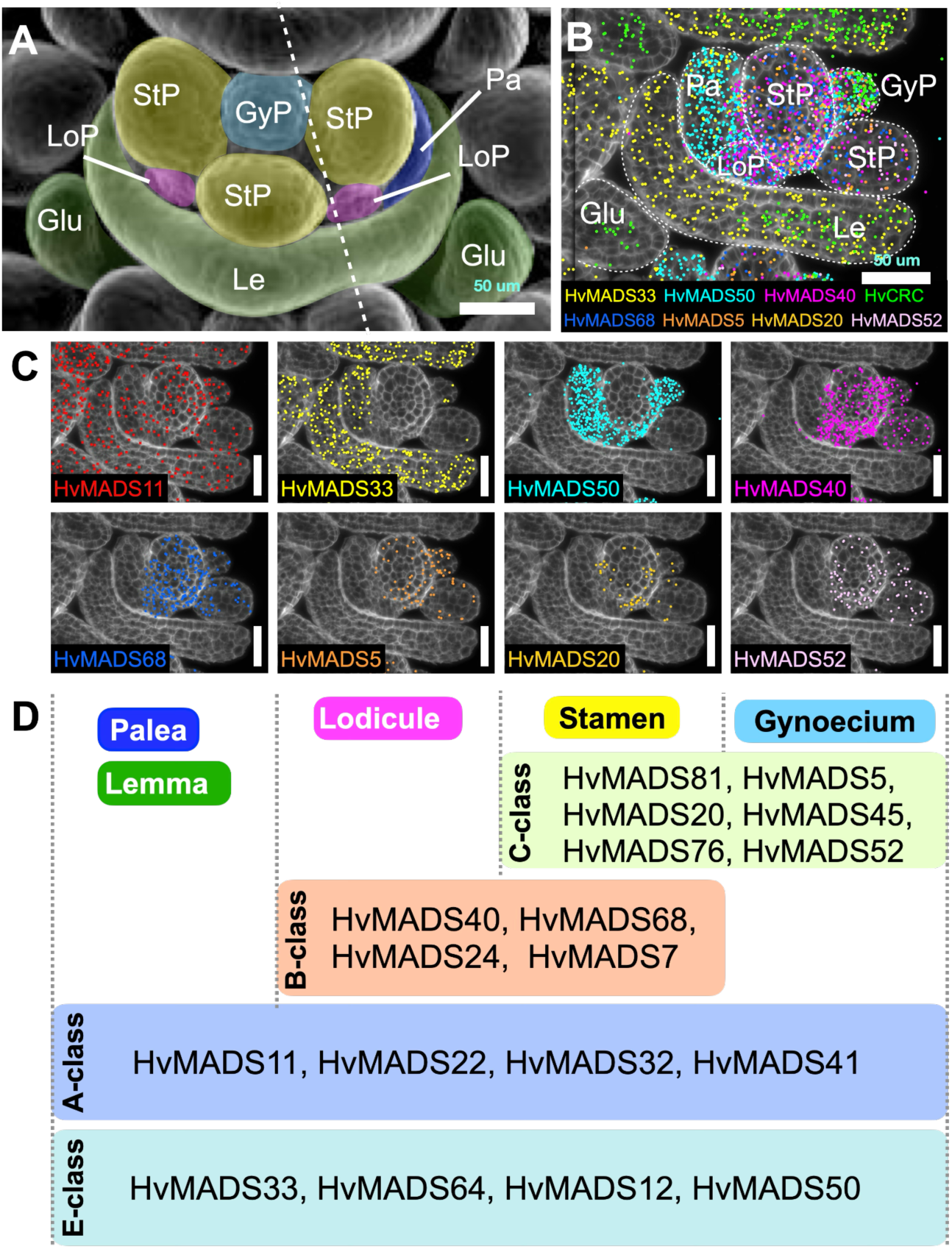
Inferred roles of *HvMADS* genes in the ABCDE model of whorl organization. (A) Scanning electron micrographs of one floret. (B) Expression profiles of *HvMADS* genes in floret organs (glumes, Glu; lemma, Le; palea, Pa; lodicule primordium, LoP; stamen primordium, StP; gynoecium primordium, GyP). (C) Individual expression profiles of selected *HvMADS* genes. (D) Putative role of MADS-box transcription factors inferred by pseudotime analysis in the ABCDE model of floret whorl formation. The dashed line in (A) represents the approximate orientation in the section in (B). Scale bars: (A)–(C) 50 µm.

### Gene expression patterns in distinct phases of meristem development

Using the *BARVISTA* virtual microdissection tool, we integrated the pT data from the scRNA-Seq dataset with our imputed gene expression dataset and identified genetic determinants that are both temporally and spatially associated with specific developmental stages of primordia. Barley spikelet primordia develop in a distichous arrangement, with a developmental gradient along the apical-basal axis. We selected the first 10 primordia on a single side of the sections (P1-P19, only odd numbers), and captured spatio-temporally defined DEGs throughout spikelet development from the TSM to FM and the formation of floral organs. Each spikelet was defined by excluding the subtending lemma primordium, characterised by the expression of *HvCRC*, and cells expressing *VRS4* were used to delimit spikelet boundaries (**Fig6A**). We then identified 637 marker genes by comparing their expression levels in each primordium with their expression in the total number of selected cells (**Fig6A, SupplTab17**). Most DEGs were detected in the early stages P1 to P9, reaching a peak of 303 DEGs in P5, which marks the transition between TSM and CSM/LSM. No DEGs were detected from P11 to P15, and the number of DEGs gradually increased from P17 (5) to P19 (150). This allowed us to subdivide spikelet development into three phases: first, SM identity is defined by transcriptional reprogramming in cells of the early spikelet primordia, which differentiate into various tissue types characteristic of the SM, becoming morphologically distinct at P11; second, rapid growth occurs during P11 to P15 without significant changes in gene expression; and third, the FM undergoes differentiation, with changes in the gene expression marking the differentiation of the floret organs (GyP and StPs) (**Fig6A**). Among the DEGs, we identified 36 TF genes expressed at specific spikelet development stages (**Fig6B, SupplTab18**). Corroborating our pT data, *HvMADS42* (5HG0511250) was upregulated in P1-P7, *HvMADS33* (4HG0396400) in P7-P11 after the TSM-to-CSM transition, and *HvMADS50* (6HG0604360) from P11 onwards, when the FM is established. Other DEGs in P5 included those encoding nuclear factor Y subunit B-10 (*HvNF-YB10*, 2HG0126410), two uncharacterised basic leucine zipper domain (bZIP) TFs (6HG0570630, 7HG0746810) and the ethylene-responsive TF *HvERF61* (1HG0090590). *HvMADS64* (7HG0654930) was significantly upregulated in the CSM (P7). Between P11 and P17-P19, we observed organ growth, but no specific marker genes were detected until floret organs started to differentiate (P17 to P19). The TF genes characterising floret development included *HvMADS7* (1HG0065060), *HvMADS20* (3HG0243770), and *HvMADS76* (*HvMADS16*, 7HG0721170). The transcriptional and morphological similarity of primordia from P11 to P15, after the prominent transcriptional reprogramming during early spikelet development, indicated the establishment at P11 of transcriptionally defined spikelet organs. To test this, we used *BARVISTA* to select cells from the adaxial part of spikelet P11 (FMad), which forms the rachilla, from the floret meristem (FM) and from the abaxial side (FMab) where the lemma is initiated (**Fig6C**, **SupplTab19**). We found two YABBY TF genes as markers for FMab: *HvCRC (*4HG0396510), the ortholog of *DROOPING LEAF (OsDL)* in rice, *DL2* in maize and *CRABS CLAW* in Arabidopsis; and *HvTOB1 (*2HG0184460), the ortholog of *ZmYABBY14*/*TOB1*/*ABNORMAL FLOWER ORGANS*, all related to leaf-like organs such as the lemma in barley ^49–54^. On the adaxial side (FMad), we found WUSCHEL-RELATED HOMEOBOX PROTEIN (*HvWOX3*, 1HG0010970), the ortholog of *LATERAL LEAF SYMMETRY*/*OsWOX3* in rice, *ZmWOX3a*/*ZmWOX3b* in maize, and *PRESSED FLOWER* in Arabidopsis, which are involved in lateral organ formation ^55–57^. Other markers included *HvMONOCULM1* (7HG0720900), the GRAS-TF gene related to *ZmGRAS43* in maize, *MONOCULM1* in rice, and *LATERAL SUPPRESSOR* in Arabidopsis, which contribute to axillary meristem formation and branching in rice and Arabidopsis ^58–60^. In the central domain (FMc), the top candidate was *HvMADS50* (6HG0604360), related to rice *OsMADS6*/*MOSAIC FLORAL ORGANS1*, maize *BEARDED EAR-1*, and Arabidopsis *AGL6*, which supports floral meristem establishment ^61–63^. We also detected *HvLEAFY* (2HG0194240) ^64^ and 59 transposon-related genes specifically expressed in the FMc.

**Fig6:**
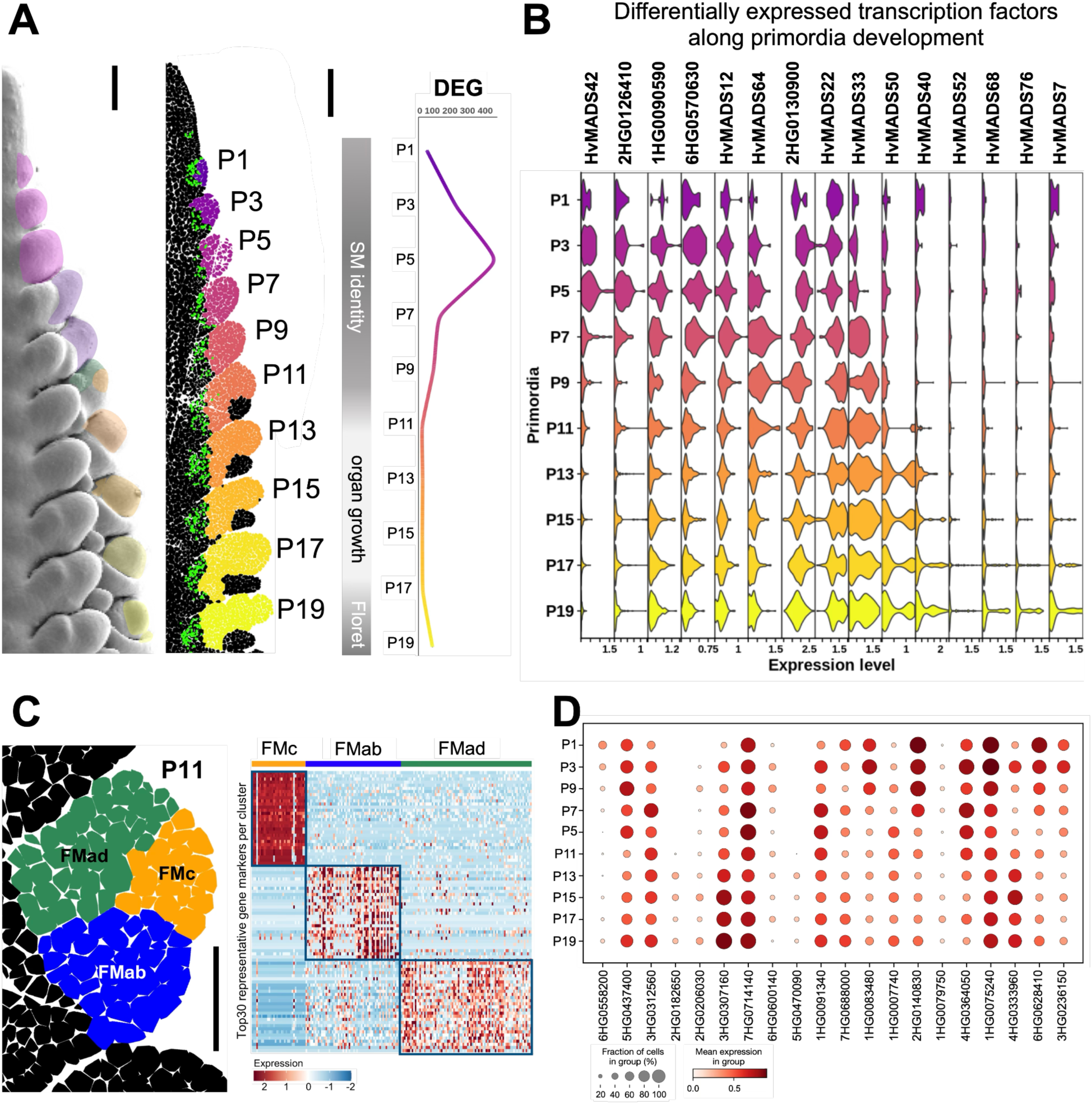
Isolation of stage- and domain-specific genetic determinants using *BARVISTA*. (A) Apical–basal gradient of spike development: formation of TSM (P1–P3), transition (P5) to CSM (P7–P9), establishment of FM (P11–P13) and floret whorl organization (P15–-P19). *VRS4* expression (green) was used to delimit primordia boundaries. P5 shows the higher diversity in imputed differentially expressed genes (DEGs). (B) Expression levels of selected DEGs encoding transcription factors across primordium development. (C) Floret meristem divided into central (FMc), adaxial (FMad) and abaxial (FMab) domains, and heat map with the top 30 DEGs for each FM region. The heat map shows selected examples of imputed DEGs using a Model-based Analysis of Single-cell Transcriptomics (MAST), with *p_adj_* < 0.01 and with enrichment for cells that express genes at double or more the level of the remaining cells (pct.1/pct.2 ≥ 2). Scale bars: (A) 100 µm; (C) 50 µm. (D) *PERSIST* identified 20 genes with specific combined expression patterns for each primordia stage.

These results show how we can use *BARVISTA* to subdivide spikelet development into different transcriptionally distinct phases and identify relevant TFs marking the differentiation of specific spikelet organs even before they become morphologically distinct. We then used PERSIST to identify gene expression combinations suitable as barcodes for stages P1-P19 ^38^ (**Fig6D, SupplTab20)**. The barcode can be used to rapidly assign a cell of unknown origin to the correct primordium, based on the similarity of gene expression to the identified barcode genes. Such a barcoding system will determine single cell identities more precisely in future experiments, estimating a cell’s position within a tissue even in the absence of spatial experimental data.

### Characterising mutant phenotypes at the single-cell level via gene expression

Spikelet identity is determined by the TFs COM1 and COM2, which act in partially independent pathways. Plants carrying mutations in both *COM1* and *COM2* develop branched inflorescences, which was interpreted to be a reversion of CSMs into indeterminate IM-like structures that generate multiple spikelets on their flanks ^9,65^ (**Fig7A,B,D,G,H,J**). Using the selected genetic markers for spikelet formation (*VRS4*, *HvCRC* and *HvLOG1*), we compared gene expression profiles in the apical and lateral meristems between WT and *com1a;com2g* inflorescences. Early expression profiles appeared identical (**Fig7C,F**), but when WT CSMs generated different spikelet organs, *com1a;com2g* CSMs grew indeterminately, and the onset of *VRS4*, *HvCRC* and *HvLOG1* expression in the *com1a;com2g* background was reduced and delayed (**Fig7E,I**; white asterisks), resembling WT inflorescence tips. We integrated smRNA-FISH data from WT and *com1a;com2g* spikes and identified cell-type specific clusters based on similar gene expression profiles, which were mapped onto the corresponding spike section. At the gene expression level, WT SMs gradually acquired FM and floret identities, whereas cells in *com1a;com2g* branches maintained tip-like identities (**Fig7K**).

**Fig7.**
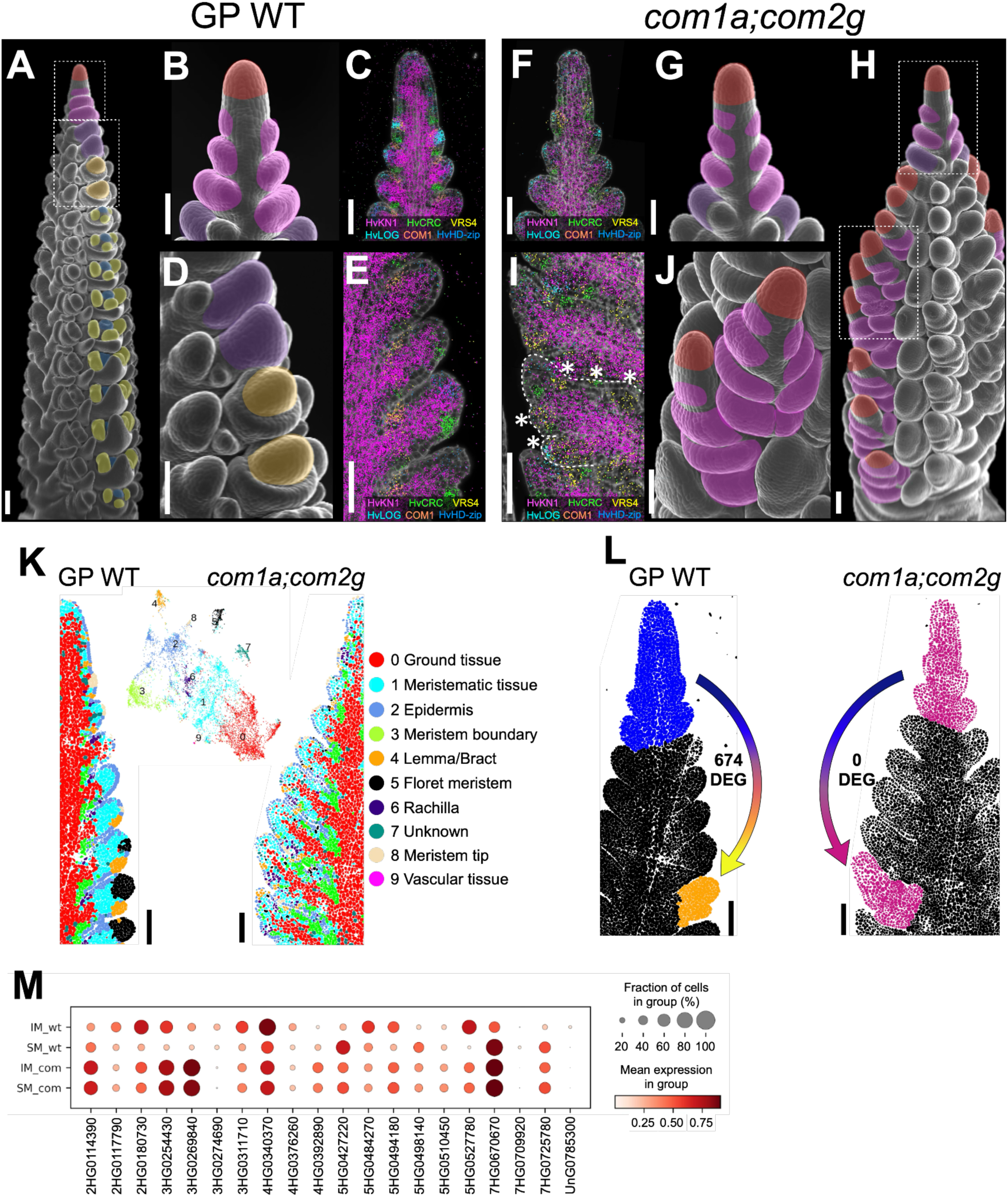
Imputed differential gene expression for molecular phenotyping at the single-cell level. (A) Scanning electron micrograph (SEM) of barley spike indicating the IM (red), TSM (pink), CSM (orange), stamen primordia (yellow) and gynoecium primordia (blue). (B) Close-up of one developing spike highlighting TSM formation, equivalent to the upper area dashed in (A). (C) Gene expression patterns associated with the establishment of TSM. (D) Close-up from (A) showing the CSM and its transition to FM. (E) Marker gene expression profiles during the transition from TSM to FM. (F) Marker genes expressed in the upper part of a *com1a;com2g* developing spike. (G) SEM of an equivalent region as shown in (F). (H) Developing spike of *com1a;com2g;* initiation of TSM (upper dashed rectangle) and further IM-like development (lower dashed rectangle). (I) Expression profile of marker genes in an IM-like branching structure, with asterisk indicating the formation of ectopic SMs. (J) SEM of branched structure equivalent to the dashed (lower) region in (H). (K) Spatial clusters of smRNA-FISH data from WT and *com1a;com2g* after integration. The WT SMs acquired FM and floret identities (cluster 5 FM, black cells). In *com1a;com2g*, cells with FM identity (black, cluster 5) are underrepresented and branches maintained tip-like identities. (L) Imputed differentially expressed genes between the upper part of the spike, including the IM and the first four primordia (P1–P4, blue), and P13 (FM, orange) in WT and its counterpart in *com1a;com2g*. (M) Gene expression barcodes for SM and IM of WT and *com1a;com2*g (as selected in L). The barcodes show expression levels of 20 genes that distinguish between the tissues. Genes were selected using *PERSIST*, based solely on their expression patterns. Scale bars: 100 µm.

We identified transcriptional signatures underlying spikelet determination by imputing gene expression differences between the inflorescence tips, spikelets and the *com1a;com2g* IM-like branches. The IM-like branch bears four axillary SMs, so we selected the inflorescence tip including the first four spikelet primordia (**Fig7L**). In WT plants, we identified 674 DEGs between the inflorescence tip and determinate spikelet, whereas *com1a;com2g* branches showed no DEGs in comparison to the inflorescence tip (**SupplTab21**). This demonstrates how specific transcriptional signatures can be used to describe and define phenotypic profiles, independent of morphological comparisons. To minimise the number of gene expression profiles needed to characterise a specific organ, we used *PERSIST* to generate gene expression barcodes for WT and mutant inflorescence tips and AMs. Expression barcodes for the WT inflorescence tip were clearly distinct from those of the determinate spikelet, whereas those of the *com1a;com2g* inflorescence tip and branches were indistinguishable. Interestingly, expression barcodes for the mutant inflorescence tip and branch shared commonalities with both the WT inflorescence tip and WT spikelet, indicating that both the *com1a;com2g* inflorescence tip and the branches represent an intermediate state. Comparison of MADS-TF gene expression revealed that *COM1* and *COM2* promote *HvMADS33* and *HvMADS64* expression from P7 onwards, and *HvMADS20* and *HvMADS50* expression in FMc from P11 onwards (**SupplFig16**).

Complete gene expression profiling together with the generation of tissue/stage-specific expression barcodes thus facilitates and simplifies not only the understanding of mutant phenotypes and their underlying changes in gene expression, but also the identification of cell and tissue identities in complex genetic backgrounds.

## Discussion

### Extending spatial gene expression datasets via gene expression imputation

The location of cells in plant tissues is generally fixed. Most cells are only passively displaced during growth while keeping contact with their neighbours, establishing local clones. Positional information is imparted by the perception of juxtacrine or symplasmic signals from direct neighbours, and/or by exposure to specific concentrations of morphogens or abiotic factors such as light. Such spatial inputs modulate the same gene expression network operating in different cells and can generate position-dependent outputs that should be taken into account when interpreting phenotypes from gene-network perturbations. However, methods for the spatial analysis of gene expression cannot yet accommodate the full complexity of transcripts in a cell. For probe-based methods, this reflects the molecular crowding of complex and bulky probes, which are needed to increase the sensitivity or specificity of target RNA detection. Methods based on cDNA sequencing are limited by the efficacy of RNA capture from tissues. Both methods generally detect transcripts from fewer than 5,000 genes, and information on expression dynamics is limited. Alternatively, scRNA-Seq reveals the expression dynamics of many thousands of genes, but lacks spatial information. Our approach combines the benefits of both methods to address the problem of space in gene expression studies ^26^. We obtained cell-resolved, quantitative data on gene expression during barley meristem development, covering a maximum number of genes, by combining scRNA-Seq and cell-resolved transcript detection on tissue sections. Using a small set of probes as anchors to reliably detect transcripts at many different expression levels, we were able to match cells from tissue sections with those from the scRNA-Seq dataset, using CS followed by the calculation of a weighted average from the most similar cells for each gene. We then used gene expression information from the scRNA-Seq dataset to impute the missing expression values of almost all barley genes for the cells in tissue sections. Validation tests showed that the imputed gene expression values were highly reliable, with the exception of protoplasting-induced genes. We focused on key imputation parameters to identify our method’s limitations, and found that the 81 anchor genes from the smRNA-FISH experiment were able to identify similar cells between the datasets, but the selection of anchor genes could be optimised using the deep learning tool *PERSIST* for gene selection. We found that for a dataset as complex as the barley inflorescence, scRNA-SEQ data from 10,000 cells and 20 anchor genes were sufficient for gene expression imputation, and to identify all cells within a tissue. Gene combinations can be used to differentiate between cell populations, and to establish simplified gene expression barcodes for specific cell identities or states in a dataset, revealing complex changes in gene expression, in mutants or during developmental progression. Future applications could integrate additional data types, such as proteomic or metabolomic data. Within the given experimental limitations, our imputation approach enables gene expression analysis by virtual microdissection of individual cells for the discovery of cell or location specific characteristics, or to determine functional domains within meristems that underlie specific traits.

### Dynamic gene expression underlying the formation and specification of primordia

*KN1* and its orthologs promote meristem fate in many cereal species, and we found that the downregulation of *HvKN1*, combined with *HvHD-zip* expression, is a hallmark of primordia initiation in the vSAM and IM. Leaf founder cells were predominantly marked by their expression of *HvCRC*, which was also expressed in spikelet founder cells, but polarised to the abaxial side, and marked cells that would form the suppressed bract. On the adaxial side, in addition to *HvLOG1*, we observed the specific expression of *VRS4* and *COM1*, encoding TFs that promote spikelet determinacy in barley.

*KN1* mRNA in meristems was found to be restricted to corpus tissue, and was only sporadically found in the tunica layers in maize or rice. We have shown here that *HvKN1* is expressed in the corpus of spikes and, at a lower level, of vSAMs. *HvKN1-positive* cells from the corpus of vSAM, and *HvKN1-negative* tunica cells from spikes showed positive correlation in gene expression, indicating shared developmental paths (**SupplFig10E,F**). *HvKN1-positive* tunica cells are located at the tips of IMs and SMs, whereas tunica cells in leaf primordia tips of the vSAM lacked *HvKN1* expression. The expression of *HvKN1* is strongly correlated with *HvLOG1*, which promotes cytokinin synthesis. Such a cytokinin source in meristem tip tunica cells (but not those in the tips of leaf primordia), generated via the *HvKN1*/*HvLOG1* gene network, could establish the *WUS/WOX* expression domains in the IM and newly formed SMs ^66^ and provide a feedback signal that maintains *HvKN1* expression, thereby acting as a local organiser that promotes meristem maintenance from the outermost cell layer (**Fig8A,B**).

**Fig8.**
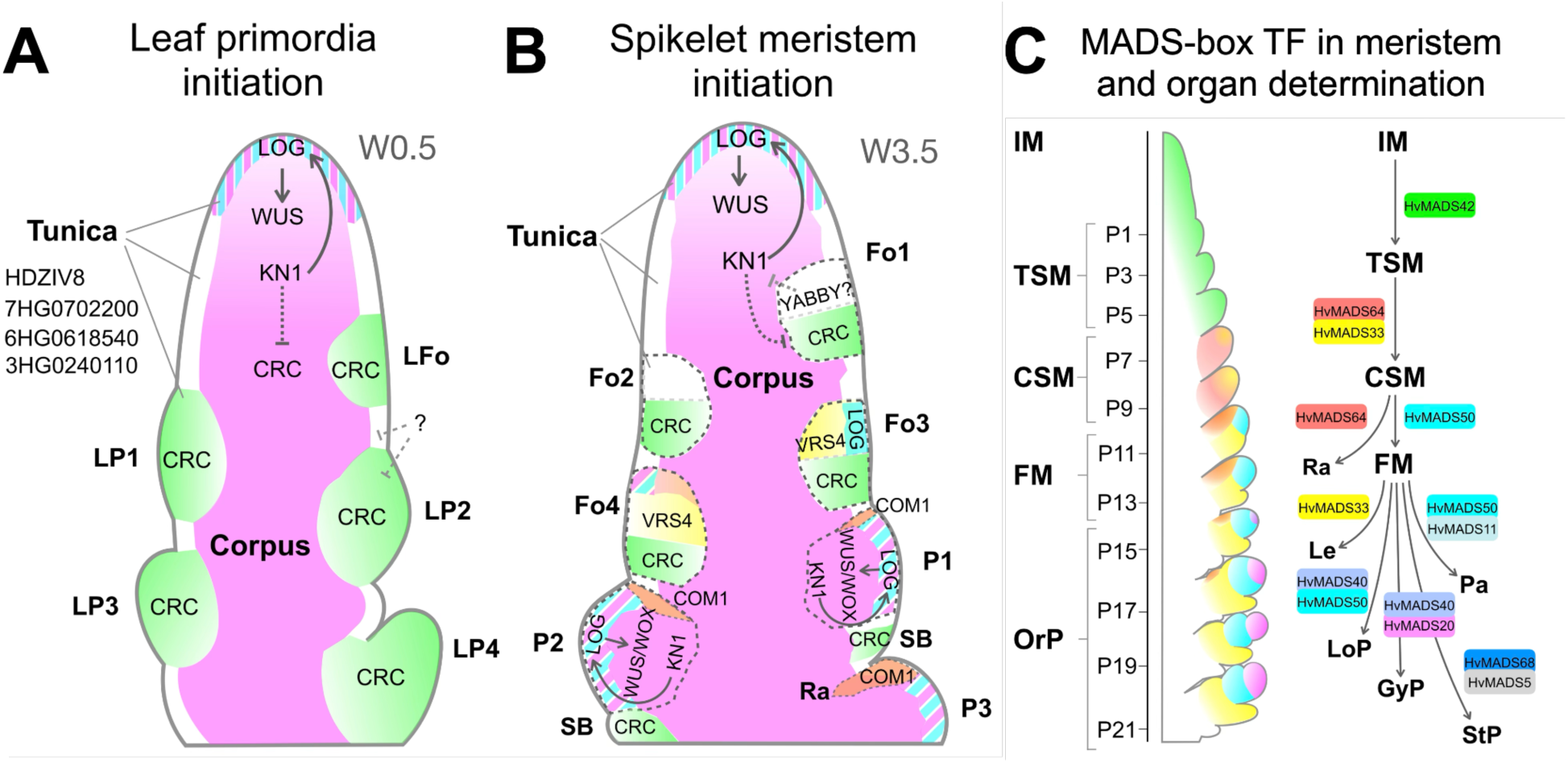
Model for primordia initiation and meristem organisation in vegetative SAM and developing inflorescences. (A) Model for leaf primordia specification and initiation in vegetative SAM. HvKN1 (magenta) is expressed in the corpus and the tip of the SAM; HvKN1 positively regulates LOG (magenta and cyan stripe pattern), which promotes cytokinin biosynthesis. Cytokinin upregulates HvWUS/HvWOX genes to organize meristem function. Leaf primordia are specified in founder cells (LFo) depleted for HvKN1, with HvCRC (green) as the earliest sign of leaf primordia determination (LP). (B) Inflorescences initiate triple spikelet meristems (TSM). Founder cells (Fo1-Fo2) lack HvKN1 expression; HvCRC (green) in the basal domain marks Fo cells of the suppressed bract (SB), YABBY TFs become expressed in the apical domain. At Fo3-Fo4, VRS4 (yellow) and LOG (cyan) are expressed in the apical domain, possibly specifying TSM identity. From P1 onwards, HvKN1 expression is restored and extends into the tunica. LOG expression fosters cytokinin accumulation, which upregulates HvWUS/HvWOX genes to organize and pattern the newly formed TSM. Expression of COM genes (orange) then determines the rachilla meristem (Ra). (C) MADS-TFs are major regulators in meristem and floret organ determination. HvMADS42 (green) is expressed during the transition from IM to TSM. HvMADS64 and HvMADS33 (red and yellow respectively) are expressed in CSM, then HvMADS64 specifically in Ra. HvMADS50 (cyan) is specific for the central zone of FM, HvMADS33 for the lemma primordia (Le), the first floral organ formed. Combinations of HvMADS-TF genes mark the specification of other floral organs (OrP). HvMADS50 (cyan) and HvMADS11 are expressed in the palea (Pa); HvMADS50 and HvMADS40 in the lodicule primordium (LoP); HvMADS40 and HvMADS20 in the gynoecium primordia (GyP) and HvMADS68 and HvMADS5 in stamen primordia (StP).

Virtual microdissection using *BARVISTA* allowed us to find specific marker genes for the IM, TSM founder cells and TSM primordia (**Fig3F-H**). These included genes encoding the flowering regulators HvFT2 and HvFD7 for IM, the TFs HvTOB1 and HvTOB2 for Fo cells, and HvSCR1 for P1 or TSM. In maize, stem cells in the vSAM and IM, marked by *ZmWUS1* and *ZmCLE7* expression, are often difficult to recover from protoplast preparations ^22^. Our scRNA-Seq dataset included a significant number of cells expressing *HvWUS1* and *HvCLV3*. Furthermore, maize *PYRUVATE ORTHOPHOSPHATE DIKINASE* (*ZmPPDK*) is highly co-expressed with *ZmCLE7* in the IM ^22^, and we found *HvPPDK* (1HG0062940) transcripts enriched in the barley IM, indicating that expression imputation can digitally reconstruct the IM and facilitate marker gene identification from limited tissues. Finding marker genes specific for the IM is challenging because many genes relevant for IM development are also expressed in SMs, including *HvKN1*, *HvLOG1* and *HvHD-zip*/*HvHDZIV2*/*HvOCL2* (**Fig3G**) or *HvMADS42*/*MADS34* and *HvMADS64*/*MADS5* (**SupplFig14**). Alternatively, the expression dynamics of several genes can be combined into highly specific expression profiles, using a barcoding strategy, to identify specific cell types or tissues.

### Subclustering and pseudotime analysis predict meristem transitions and the path to floret organ establishment

Barley spikes at W3.5 comprise all major stages and transitions in the developmental path towards floret formation, and we used *COM1* expression in the TSM as a starting point for pseudotime analysis. Double mutants of *COM1* and *COM2* develop branched spikes, suggesting a defect in the early stages of CSM determination ^9^. *VRS4* is already expressed in TSM Fo cells (**Fig3G**) and follows a pseudotime expression profile similar to *COM1* (**FigSuppl13B,C**). Following the developmental trajectory by pseudotime analysis, we identified genes marking the major transitions from IM to floret organs in the inner whorl. Genes modulated during the transitions from IM to FM (up to pT9) showed a higher diversity in patterns of gene modulation. Later in development, from FM to the formation of floret organs, patterns were less diverse, but more genes were modulated (**Fig4E, SupplTable16**). We then reconciled pseudotime and canonical time using the plastochron and developmental time based on meristem or organ formation (**Fig4F**). MADS-TFs are master regulators of floret organ development, and by using their pseudotime modulation, we assigned them to the ABCE model of floret development (**Fig5**). Virtual microdissection with *BARVISTA* corroborated the pseudotime analysis data and identified potential regulators of key developmental transitions. Many DEGs were detected at stage P5, when the TSM separates into a determinate CSM and LSMs. These DEGs included *HvNF-YB10*, *HvERF61*, *HvMADS33* and *HvMADS42*. NF-Y subunits, orthologs of barley *HvNF-YB10*, mediate epigenetic control of the floral transition in Arabidopsis ^67^, and an *HvERF61* ortholog regulates the production of key floral constituents in *Osmanthus fragrans* ^68^. Interestingly, the mutation of *HvMADS33*/*MADS1* generated branched spikes at high temperatures ^11^, and the mutation of the closest ortholog of *HvMADS42* in *Setaria italica* (*SiMADS34*) generated panicle-like inflorescences with long primary branches ^69^. The rice ortholog of *HvMADS42* (*OsMADS34)* cooperates with *OsMADS5* to promote meristem determinacy, and *osmads5;osmads34* double mutants generate secondary, tertiary or even quaternary branches ^70^. *HvMADS64*, an ortholog of *OsMADS5*, was significantly upregulated in the WT CSM (P7), whereas the branched inflorescences of *com1;com2* mutants failed to activate the expression of *HvMADS20*, *HvMADS33*, *HvMADS50* and *HvMADS64*. We conclude that a key function of TFs expressed from P5 onwards is specification of CSM identity and branch suppression. *HvMADS7* (ortholog of Arabidopsis *PISTILLATA*), *HvMADS20* (ortholog of rice *OsMADS3*) and *HvMADS76*, which regulate petal, carpel and stamen development in Arabidopsis, rice and barley ^71–73^, are probably stage-specific master regulators of barley floret organ development (**Fig8C**).

We have showcased how the computational integration of scRNA-Seq and spatial transcriptomics data significantly advances our understanding of plant meristem development and organ formation, allowing us to identify and study the underlying gene regulatory network. The gene expression imputation approach that we developed closes a very important gap in our understanding of developmental processes, by allowing us to determine or deduce quantitative expression data for all genes with cellular resolution. We also found that the high costs currently associated with spatial and single cell transcriptomics can be reduced using imputation approaches, aiming to generate multidimensional atlases for developmental and physiological studies in plants. *BARVISTA* allows the intuitive navigation of these data to identify gene expression patterns of specific genes, individual cells or cell groups with high spatial resolution. Combined with functional dissection of promoter elements, we should soon be able to redesign cereal inflorescence architectures and adapt crop plants to new challenging environments. *BARVISTA* currently offers only a snapshot of barley development, with expression data for specific developmental time points limited to two dimensions. Extending this approach to 4D is now within reach.

## Data availability

Metadata associated with this investigation can be accessed here: http://purl.org/barvista/arc. BARVISTA is accessible here: http://purl.org/barvista/home.

## Acknowledgements

Research in the R.S. lab and M.v.K laboratories is funded by the Deutsche Forschungsgemeinschaft (*CEPLAS*, EXC2048, and *CSCS*, FOR5235). B.U. acknowledges funding through CEPLAS EXC2048 p.no. 390686111 and the NFDI DataPlant NFDI p.no. 442077441.

## Methods

### Plant Growth Conditions

WT tissue was collected from *Hordeum vulgare* cv. Golden Promise plants grown in Minitray soil (Einheitserde, Einheitserde Werkverband) in 96-well trays with 4g/L Osmocote Exact Hi.End 3-4M, 4th generation (ICL Group) under long day (LD) conditions (16-h photoperiod, 20/16 °C day/night temperature). Before seed sowing, grains were pre-germinated for 4 days at 4 °C in Petri dishes lined with wet paper. Golden Promise takes ∼30 days after emergence to reach W3.5 under these conditions. For the scRNA-Seq work, 25 biological replicates per sample were harvested and pooled and the overall experiment was carried out three times. The *com1a;com2g* mutants were grown under the same conditions and collected when WT (Bowman) reached W3.5. We dissected ∼80 plants for vSAM library preparation 11 days after sowing in soil. The tissue was collected for protoplast isolation before verification at the W0 stage.

### Protoplast isolation

Protoplasts were generated from developing inflorescences (W3.5, when the stamen primordia emerged). The tissue was directly collected in six-well plates containing 3 ml Washing Solution I (WSI; 0.4 M mannitol, 20 mM MES, 20 mM KCl, 10 mM CaCl_2_, 0.1% BSA, pH 5.7). For cell wall digestion, the tissue was chopped in WSI using a scalpel and immediately digested for 140 min in 4 ml enzyme solution (0.4 M mannitol, 1.2% Cellulase R10, 1.2% Cellulase RS, 0.36% Pectolyase Y-23, 0.4% Macerozyme R-10 (all from Duchefa), 20 mM MES, 20 mM KCl, pH 5.7, 10 mM CaCl_2_, 0.1% BSA, 0.06% β-mercaptoethanol). Protoplasts were then passed through a 40-µm cell strainer (Scienceware Flowmi) into microfuge tubes for centrifugation (250 *g*, 2.5 min, 4 °C). The pellets were gently washed once with WSI and twice with WSII (0.6 M mannitol, 2 mM MES, 0.1% BSA, pH 5.7) with centrifugation under the ame conditions described above between each wash. The protoplasts were passed through the 40-μm cell strainer and stained with calcein AM (10 μM) and DRAQ7 (1.5 μM) for 5 min at room temperature. An aliquot was loaded into a hemocytometer to score cell viability using a BD Rhapsody scanner. Only samples exceeding 70% viability were loaded into the BD Rhapsody Express system for partitioning and mRNA capture.

### scRNA-Seq

We loaded ∼20,000 cells per replicate. Single-cell libraries were prepared using the mRNA Whole Transcriptome Analysis (WTA) kit (BD-Biosciences) and libraries were sequenced on the Illumina NextSeq2000 platform, aiming for 50,000 reads per cell. Raw scRNA-Seq data were analyzed using the BD WTA Rhapsody Analysis Pipeline. Gene reads were aligned to the *H. vulgare* cv. Morex V3 reference genome (https://doi.org/10.5447/ipk/2021/3).

### Subtraction of DEGs induced during protoplast isolation

Total RNA was extracted from three replicates of fresh protoplasts and intact inflorescences using the Direct-zol RNA Miniprep Plus kit (Zymo Research), followed by bulk RNA sequencing using the Illumina NextSeq platform (Biomarker Technologies). Reads were aligned to the *H. vulgare* cv. *Morex* V3 reference genome and converted to transcripts per million (TPM) by adapting a custom Python script (https://github.com/cvanges/spike_development). DEGs were defined using a count-based Fisher’s exact test in the *R* package EdgeR v3.32.1. The FDR was adjusted using the Benjamini-Hochberg procedure. Genes were considered differentially expressed when exceeding the threshold LogFC ≥ 3 or ≤ –3 and a FDR < 1×10^−5^. These 3,684 genes were removed from the scRNA-Seq expression matrix before identifying the most variable features for cluster analysis.

### UMAP and replicate integration

Replicates were merged using the *Seurat* package v4.0 ^36^. Low-quality cells were removed with a threshold of 800 < genes < 9,000, and cells were log-normalized. Protoplasting-affected genes were removed from the merged object, and the 7,500 most variable features were identified using the *vst* function in *Seurat*. Next, *FindIntegrationAnchors* identified anchors to integrate the three replicate datasets using 30 dimensions. The integrated expression matrix was retrieved using *IntegrateData*. Dimensionality was reduced by scaling the expression matrix using *ScaleData,* and then by principal component analysis (PCA). The top 50 components were selected for further analysis. Cells were clustered using a *k*-nearest neighbors and SNN graph method. *FindNeighbors* and *FindClusters* were applied with a resolution of 1.3. Dimensionality reduction was then achieved using the UMAP algorithm with the top 35 principal components and a minimal distance of 0.01.

### Cluster-specific gene marker identification

Genes enriched in each cluster were identified using the *FindAllmarkers* function in *Seurat* with thresholds of: Log.FC = 0.25, pct1 >0.25, pct2<0.3, P_adj_ < 0.001. Differential gene expression was calculated using the function *FindAllMarkers* and *Model-based Analysis of Single-cell Transcriptomics* (*MAST*) ^74^.

### Multiplex smRNA-FISH

Developing inflorescences were fixed in phosphate buffered saline (PBS) containing 4% paraformaldehyde (PFA) and 0.03% TritonX-100 (Sigma) before dehydration through an ethanol series, and progressive embedding under vacuum in Paraplast X-tra (Leica). We prepared 10 μm sections on glass slides (Resolve Biosciences), which were deparaffinized, gradually rehydrated and permeabilized with proteinase K (Thermo Fisher Scientific). After re-fixation, acetylation and dehydration through an ethanol series, the slides were mounted with SlowFade-Gold Antifade reagent (Invitrogen). Probe hybridization and imaging were carried out at Resolve Biosciences using in-house protocols. The target genes are listed in **SupplTab5**.

### Pseudotime analysis

The *Seurat* object with the integrated cells was used for pseudotime trajectory analysis with *monocle3* v1.3.7 ^75–77^ in *RStudio*. Only clusters from corpus cells were analysed in detail. These were first subclustered with *Seurat* using 50 components, a resolution of 2 and a dimensionality of 35. *COMPOSITUM1* (5HG0479720) was selected as the starting point for developmental time because COM1 is specifically expressed at the branching point of developmental trajectories towards tunica, corpus or vascular fates. The subclusters were ordered according to the learnt trajectory and pseudotime-modulated genes were calculated using *graph_test* with threshold values of *Moran’s I* > 0.1 and *q* =< 0.01. Selected examples (*q_value* == 0) were visualised using *ComplexHeatmap* v2.18.0 ^78^.

### Cell segmentation

To calculate the gene expression matrix for the MC data, the DAPI image was segmented using a workflow provided by Resolve Biosciences (https://my.resolvebiosciences.com/help/segmentation). Briefly, the pipeline first runs *MindaGap* (https://github.com/ViriatoII/MindaGap) with default settings to improve the borders of the image tiles, then *Cellpose* ^79^ with the parameters model=cyto, diameter=50.0 and flow-thresh=0.5. Genes with >=70% transcripts inside DAPI segments were considered nucleus specific, and those with <=30% transcripts inside DAPI segments were considered cytoplasm specific. Finally, *Baysor* ^80^ is run using the DAPI *roi* files as prior with the parameters --no-ncv-estimation --force-2d --n-clusters=1 --prior-segmentation-confidence 0.9 -m 3, as well as the previously determined list of nucleus/cytoplasm specific genes for --nuclei-genes and --cyto-genes. The resulting MC count matrix was used for subsequent analysis.

### Integrated clustering

*Seurat* v4.3.0 was fed with count matrices from the MC and scRNA-Seq as inputs. Data quality was based on the number of cells per sample, the number of UMIs per cell, the number of genes per cell and the complexity of all four datasets. Subsequently, all genes not recorded in the MC dataset, as well as those affected by protoplasting, were removed from all count matrices, resulting in a total of 81 *anchor genes* for the IM, 76 for the vSAM, and 83 for the *com1;com2* mutant. The datasets were log-normalised based on the natural logarithm with a scale factor of 1 using *NormalizeData.* All four *Seurat* objects were merged into one using *Seurat’s merge*, and then split into a list object with *SplitObject*, with one list element containing one sample. A second normalisation step was applied across all four datasets for further harmonisation. All available genes were selected for subsequent integration using *FindVariableFeatures* and *SelectIntegrationAnchors*, with the identified integration features as anchor features, and the MC dataset as a reference to integrate the other three datasets. For the clustering of the vSAM, we tested different parameter settings in *SelectIntegrationAnchors* (**SupplTab22**). We varied *k.anchor* between 3 and 10, *k.filter* between 25 and 400, and *k.score* 10 and 50. For the final clustering of the vSAM, we selected *k.anchor=4*, *k.filter=25* and *k.score=30*, whereas default parameters were used for the IM. The datasets were integrated using *IntegrateData*, scaled using *ScaleData*, and PCA was performed using *RunPCA* with *VariableFeatures* as PCA features and 50 dimensions. Neighboring cells were calculated using *FindNeighbors* on the PCA-reduced dataset with the first 35 dimensions, and clusters were determined using *FindClusters* with varying resolutions between 0.1 and 2.0 (step = 0.1). The *com1;com2* mutant was clustered with a resolution of 0.5. The clusters were visualized based on *Seurat’s RunUMAP* results, using the PCA reduction method and 35 UMAP dimensions. The differential gene expression of each cluster was compared to all other clusters using *FoldChange* in *Seurat* v5.0.3 for each resolution (a different *Seurat* version was used due to a known error in the fold change calculation of *Seurat* v4.3.0). All genes from the reference genome were annotated using *Mercator* v4.6 ^81^. Clusters were identified using known marker genes as well as the *Mercator* annotations.

### Imputation of genome-wide gene expression

For all genes in the scRNA-Seq dataset (reference data) but not measured in the MC experiment (test data), gene expression was imputed in a two-step procedure, loosely as previously described ^82^. All raw counts were normalized to counts per million (CPM) without additional integration. The first step of imputation (cell similarity determination) required the similarity between each MC cell and scRNA-Seq cells (SCs) to be determined. We calculated the CS between each MC cell and all segmented cells within the same cluster based on the 81 previously determined anchor genes, and the result was shifted by 1 into the positive range. For the vSAM, the CS to all segmented cells within all clusters was calculated instead due to the lower number of overall segmented cells. During the second imputation step, the expression of each MC cell was imputed by taking the weighted mean gene expression in the *n = 25* most similar segmented cells using the shifted CSs from the calculated similarity matrix as weights (Equations 1 and 2). For the vSAM, the number of top *n* cells to keep was varied between 1 and 25 during an additional parameter optimization step, ultimately using *n = 4* for the final imputation.

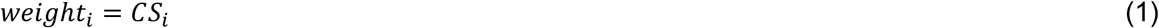

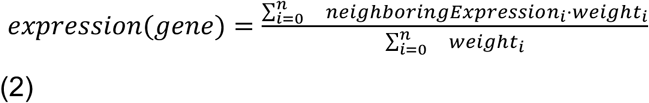

### Imputation quality assessment

To evaluate the accuracy of the imputation method, the imputed gene expression values of the MC data were compared to the CPM-normalized smRNA-FISH gene expression values. The comparison of imputed and experimentally measured values was applied for the anchor genes (81 for the IM, 76 for the vSAM, 83 for the *com1;com2* mutant) using CS as the similarity measure (Equation 3). Gene expression patterns were compared on a cell-by-cell basis. For genes not measured in the MC data, the scRNA-Seq data were used instead.

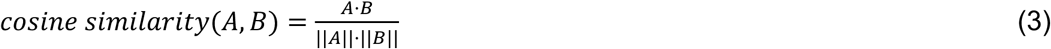

where A is the gene expression vector of the experimentally measured values, and B is the gene expression vector of the computationally imputed values.

To further evaluate the accuracy of the method, the scRNA-Seq data from the inflorescence was split into test and reference datasets using the third scRNA-Seq batch containing 5,910 cells as test cells to impute gene expression values for, and the cells of the first and second batches to impute the gene expression from. The first test set contained the same 81 genes from the MC, and the second test set contained all 48,904 genes from the scRNA-Seq experiment. For both test sets, the cell similarity determination step of the imputation was performed using the 81 anchor genes. During validation of the imputation for the first test set, we used the 81 anchor genes (scSub IM 81 genes), and for the second test set we used all 48,904 genes (scSub IM 48,904 genes).

### Imputation quality simulations

Gene expression imputation quality was simulated on different subsets of the scRNA-Seq dataset from inflorescences to investigate the impact of different settings on the imputation quality. The full dataset consists of ∼16,500 cells and 48,904 genes. Five different simulations were performed, each with five repetitions per parameter setting. Test datasets were generated containing 1,000 cells and the same genes as the reference datasets. Gene expression was validated using all 48,904 genes in the cell-by-cell comparison of imputed to experimentally measured gene expression values.

Simulation 1: Impact of reference cell number, subsets with 100–15,000 cells were tested, gene expression was imputed using 81 anchors.

Simulation 2: Impact of gene number, subsets with 5,000 cells and 20–40,000 randomly selected genes. Gene expression was imputed using the genes contained in the reference dataset during the cell similarity determination.

Simulation 3: Impact of the MC anchor genes on imputation accuracy, compared to the random gene selection from simulation 2. Reference subsets were generated in the same manner as for simulation 2 for combinations of 20–80 genes.

Simulation 4: Impact of gene selection method on imputation quality. Sets of 20–100 genes were identified using *PERSIST* {Covert, 2023 #2085; (https://github.com/iancovert/persist; initial release)} and reference subsets generated as described above. Default parameters from the provided scripts were used, with modification of the setting *flavor=‘seurat’* for the determination of highly variable genes. During unsupervised gene selection, the setting *lam_init = 0.64* was applied.

Simulation 5: Impact of gene variability on imputation quality, reference subsets were generated containing 5,000 cells and the top 250 most or least variable genes as determined using *Seurat’s FindVariableFeatures*command.

### Generation of primordia barcodes

Gene expression barcodes for different sections of the IM from WT and *com1;com2* mutant plants were generated using *PERSIST*. One barcode was generated for the sections P1–P19 and another for the meristem tip and floret meristem of the WT and *com1;com2* mutant. During file preprocessing, highly variable genes were detected using the *flavor=‘seurat’* setting. The data were split into test and training sets to run the supervised training with default settings plus *lam_init = 0.64*. After training, 20 genes were selected for each barcode using default settings.

### Data pre-processing

Multiple input files were prepared for visualization based on *Seurat* clustering results, containing cell IDs, UMAP coordinates, cluster IDs and cluster names for all calculated resolutions. A data table was prepared containing log_2_FC and p values, and annotation information for each gene from *Seurat’s FindAllMarkers* for each cluster and cluster resolution, including the top-level *Mercator* bin in the “classification” column and the leaf-level *Mercator* bin in the “annotation” column. The count matrices were CPM-normalized and rounded to zero decimal points, compressed using gzip with six compression levels, and saved separately in two matrices for the MC and SC data respectively.

Polygon coordinates were extracted in *ImageJ/Fiji* v2.9.0/1.53t (Java 1.8.0_322; https://imagej.net/ij/; access date 13 March 2024) by creating a 16-bit mask of the calcofluor image using the *roi* files from the cell segmentation. Briefly, the developed macro creates a new image file the same size as the original image with pixel values of zero, creating a black background. For each *roi* file containing one cell from the segmentation, the pixel values were changed to the cell ID, creating the 16-bit mask file. To extract polygon corner coordinates, we developed a Python script using the *pillow* library v9.4.0 (https://python-pillow.org/; access date 13 March 2024), *scipy* v1.10.1 (https://scipy.org/; access date 13 March 2024) and *scikit*-*image* v0.19.3 (https://scikit-image.org/; access date 13 March 2024), which extracts pixel coordinates for each cell and calculates their convex hull. Polygon corner coordinates and the pixel value as cell ID were saved in JSON format to create a suitable *javascript* input.

### Web-application *BARVISTA*

For data visualization, we developed a web application in *JavaScript* (http://purl.org/barvista/home) with *D3.js*v7.8.4 (https://d3js.org/; access date 26 October 2023) as the main framework. The package *dom-to-image*v2.6.0 (https://github.com/tsayen/dom-to-image/tree/master; access date 13 March 2023) was used for image downloading. The gzipped expression matrices are streamed from the server to the client, making use of *DecompressionStream* from the Compression Streams API, *ReadableStream* from the Streams API, and *TextDecoder* from the Encoding API. The web application visualizes the integrated barley data and has two main plots, one displaying the cell clusters (UMAP plot) and the other displaying the cells within tissue sections. Clusters are color-coded and identified in both plots. At the bottom, a data table displays information about a selected cluster, including the genes, their average log_2_FC values, the percentage of cells in the selected cluster that express each gene (pct.1), the percentage of cells in all other clusters that express each gene (pct.2) and their *Mercator* annotation. The table is searchable for genes or annotations of interest. The default resolution of the cluster data is 1.3, but a resolution selector allows visualization at resolutions of 0.1–2.0 to show clustering with a higher or lower number of clusters. Gene expression information can be visualized in both plots for two genes in parallel. Cells in the tissue plot can be selected with the selection tool and gene expression can be downloaded as a subset for further analysis of specific regions of interest.

### Imputed differential gene expression analysis

Subsets of cells in the tissue sections were selected using the selection cell tool in *BARVISTA*, aiming to capture IMs (IM-01 and IM-02), founder cells (Fo-01 and Fo-02), and the first properly organized TSM primordia cells (P1-01 and P1-02). The matrices containing the imputed counts were used to create a *Seurat* object and were then log-normalized. Cells from each category were considered as a cluster and the imputed DEGs were calculated using the *FindAllmarkers* function in *Seurat* and a *MAST* statistical method, with thresholds Log.FC > 0.1, pct.1 > 0.25, pct.2 < 0.5, P_adj_ < 0.05, and a pct.1/pct.2 ratio > 1.2.

## Supplementary Tables

**Supplementary Table 1.** Metrics summary for the scRNA-Seq libraries from the different replicates used in this study.

**Supplementary Table 2.** Differentially expressed genes during the protoplasting process (–3 ≤ log_2_FC ≥ 3, FDR < 1×10^−5^).

**Supplementary Table 3.** Unique markers per cluster in developing spikes at WS3.5 (pct1 > 0.25; pct2 < 0.3; p_adj_ < 0.001).

**Supplementary Table 4**. Reported marker genes used to annotate cluster/subcluster identities and their closest homologs in maize and Arabidopsis.

**Supplementary Table 5**. Genes used for multiplex fluorescence in situ hybridization.

**Supplementary Table 6.** Annotation of MADS-box transcription factor genes in barley and their putative orthologs in rice by reciprocal BLAST.

**Supplementary Table 7.** Unique markers per cluster in the vegetative SAM (pct1 > 0.25; pct2 < 0.3; p_adj_ < 0.001).

**Supplementary Table 8.** Statistical values describing the gene expression imputation results.

**Supplementary Table 9.** Gene markers for ground tissue and tunica cells positive or negative for *HvKN1* expression in developing spikes at WS3.5.

**Supplementary Table 10.** Gene markers for ground tissue and tunica cells positive or negative for *HvKN1* expression in the vSAM.

**Supplementary Table 11.** Gene markers for TSM founder and primordia, and the rest of corpus cells (ground). P_adj_ ≤ 0.05.

**Supplementary Table 12.** Imputed differentially expressed genes between IM, founder and primordial cells using MAST (p_adj_ < 0.05, pct.1/pct.2 ratio ≥ 1.2).

**Supplementary Table 13.** Unique markers per subcluster in vascular tissues (pct1 > 0.25; pct2 < 0.3; p_adj_ < 0.001).

**Supplementary Table 14.** Unique markers per subcluster in corpus tissues (pct1 > 0.25; pct2 < 0.3; p_adj_ < 0.001).

**Supplementary Table 15.** MADS-box transcription factor transcripts detected by scRNA-Seq.

**Supplementary Table 16.** Modulated genes in pseudotime analysis (Morans I > 0.1; q ≤ 0.01).

**Supplementary Table 17.** Imputed differentially expressed genes along spikelet development (P1–P19).

**Supplementary Table 18.** Imputed differentially expressed transcription factor genes marking spikelet development (P1–P19).

**Supplementary Table 19.** Imputed differentially expressed genes in cells of regions in the floret meristem central zone (FMc), abaxial (FMab) and adaxial (FMad) using MAST (p_adj_ < 0.01, pct.1/pct.2 ratio > 1).

**Supplementary Table 20.** Genes selected for the P1–P19 barcode and the WT *com1a;com2*g barcode.

**Supplementary Table 21.** Imputed differentially expressed genes between IM and FM in WT and between IM and the equivalent plastochron by position in *com1a;com2g*.

**Supplementary Table 22.** Description of anchors found during the integration of the inflorescence meristem and the vegetative shoot apical meristem, as well as imputation quality results.

