## Supplementary Figures for "Imputation integrates single-cell and spatial gene expression data to resolve transcriptional networks in barley shoot meristem development"

### Supplementary Figure1:

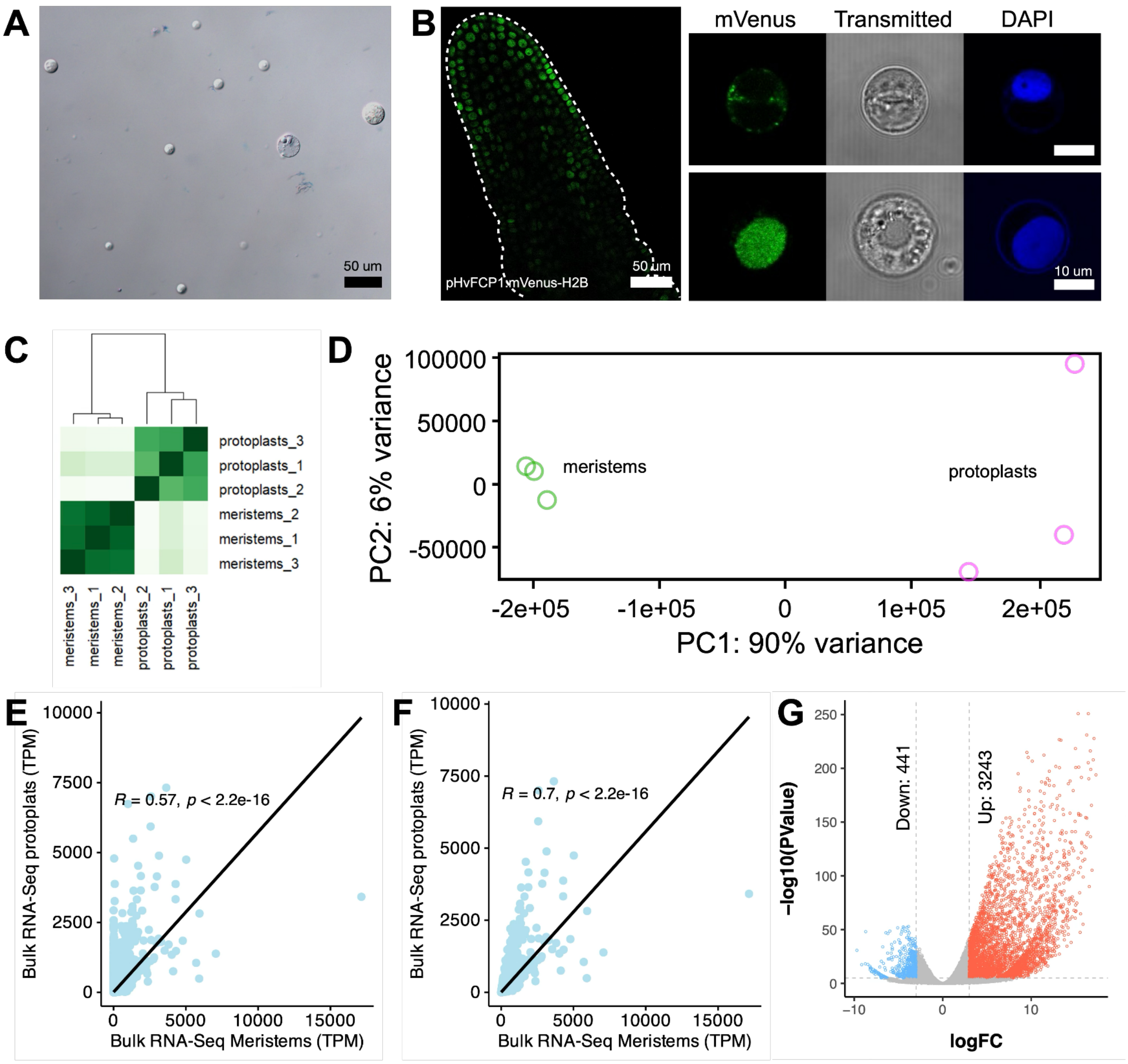


H

| Genotype | WS | Meristems | Protoplast concentration (Protoplasts/ ul) | Volume (ul) | Total Protoplasts | Digestion time (hrs:mins) |
|---|---|---|---|---|---|---|
| Golden Promise | 2.5-3.5 | 15 | 112 | 100 | 11200 | 2 |
| Golden Promise Fast | 2.5-3.5 | 25 | 56 | 500 | 28000 | 2 |
| Golden Promise | 3.5 | 20 | 60 | 600 | 36000 | 2:20 |
| Morex | 3-4 | 20 | 488 | 200 | 97600 | 2:20 |
| Bowman | 3-4 | 20 | 408 | 300 | 122400 | 2:20 |

**SuppFig1: Protoplast isolation and identification of differentially expressed genes during this process.**

(A) Size diversity of protoplasts isolated using the protocol optimised for barley spikes. (B) The transcriptional reporter line *pHvFCP1:mVenus-H2B* expressed in IM and the protoplasts recovered from this line showed the presence and absence of mVenus suggesting the isolation of diverse populations of protoplasts from the developing spike of barley. (C) Cluster correlation between replicates in bulk RNA-Seq from intact tissue and protoplasts at W3.5. (D) Principal component analysis from bulk RNA-Seq to detect differentially expressed genes during protoplasting. (E) Correlation of gene expression between bulk RNA-Seq from intact spikes (meristems) and protoplasts. (F) Correlation of gene expression after removing differentially expressed genes. In (E) and (F) Pearson's correlation coefficient: R. (G) Differentially expressed genes with $log_2FC>3$ or $log_2FC<-3$ and FDR < $1x10^{-5}$ (Supplementary Table 2). (H) Protoplast recovery with different genetic backgrounds. Scale bars: 50 µm in (A) and right panel in (B), 10 µm in left panel in (B).

### Supplementary Figure2:

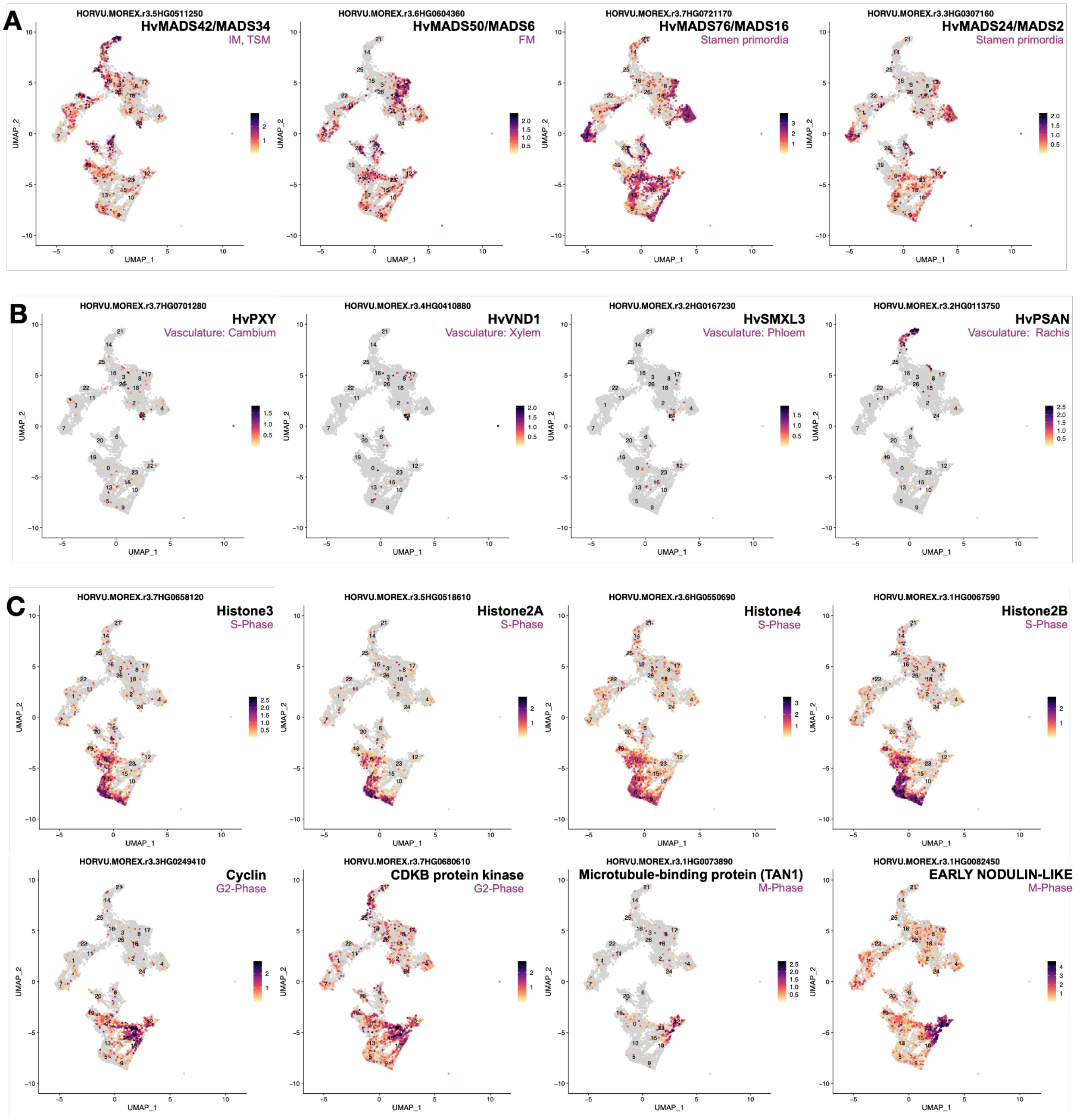

**SuppFig2: Marker genes specifically expressed in clusters.**

UMAP plots for selected marker genes associated with (A) the corpus and tunica in meristems, (B) vasculature, and (C) dividing cells (two rows with graphs).

### Supplementary Figure3:

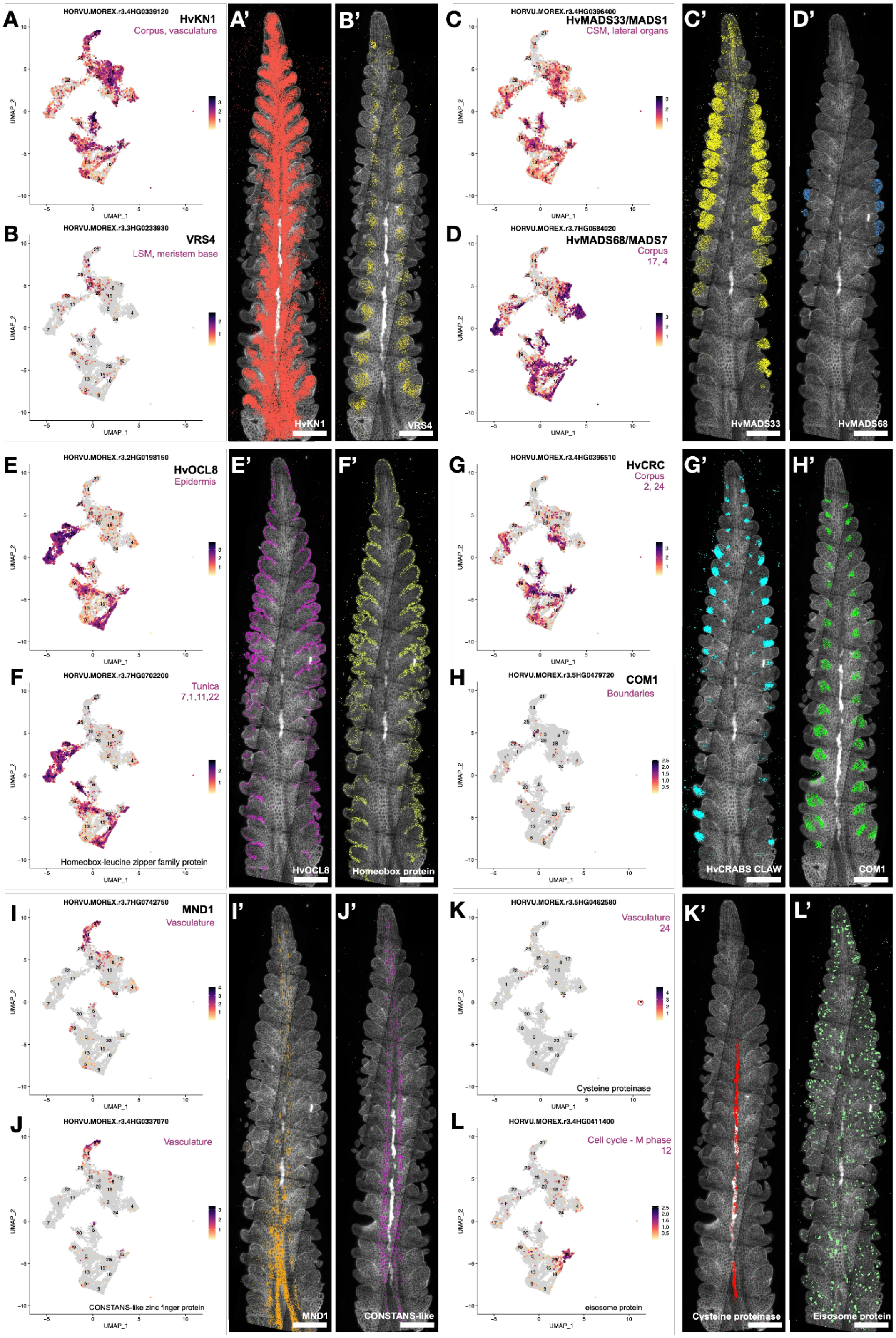

**SuppFig3: Cluster annotation and validation using known and newly discovered markers.**

(A-D) Examples of mRNA expression in scRNA-Seq UMAPs and experimental localization in developing spikes of known markers by smRNA-FISH. (E-L) Examples of mRNA expression in scRNA-Seq UMAPs and experimental localization in developing spikes by smRNA-FISH (markers newly identified by cluster analysis). Scale bars: 200 µm.

#### Supplementary Figure4:

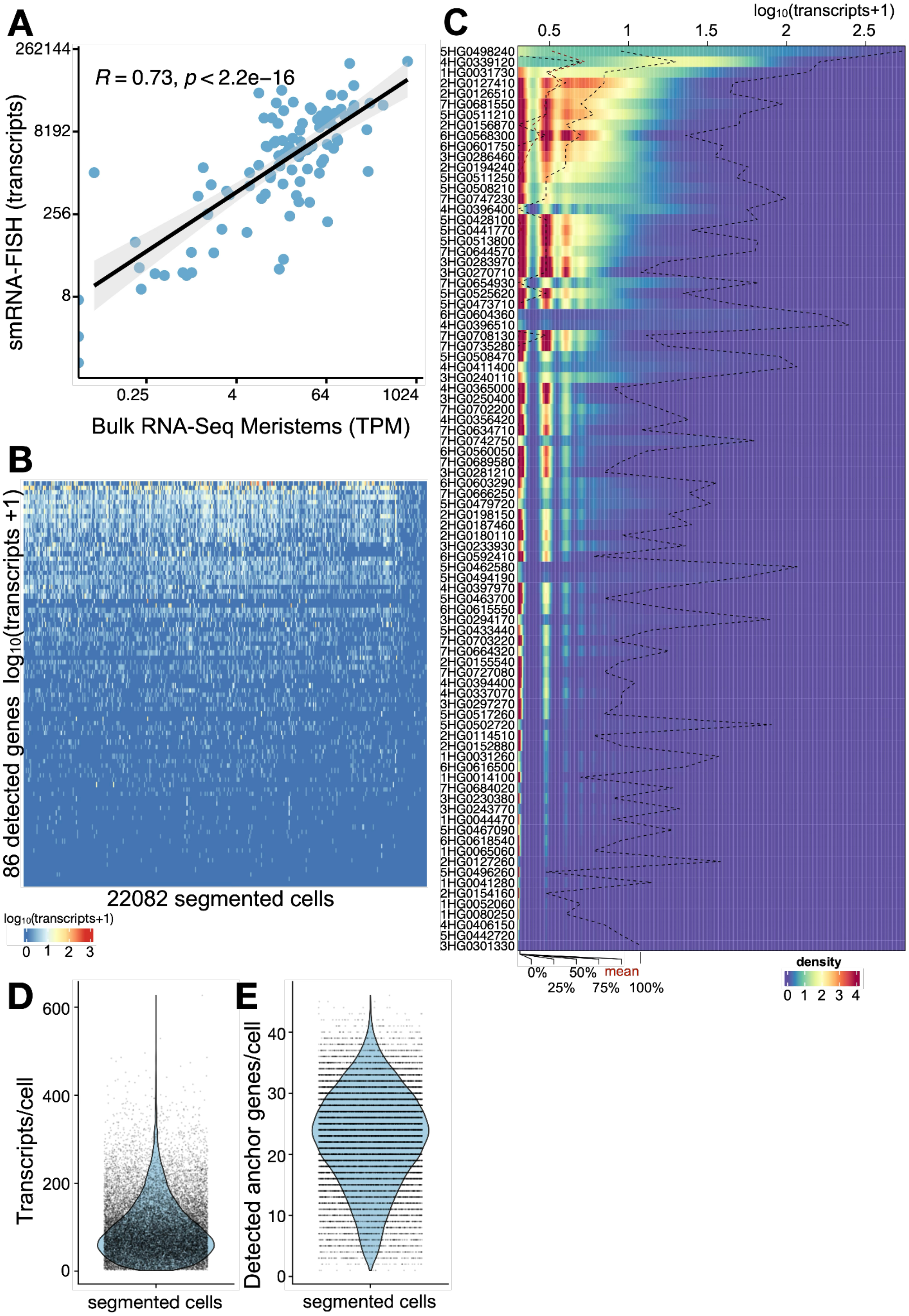

**SuppFig4: smRNA-FISH is quantitative and correlates with detected gene expression.**

(A) smRNA-FISH transcripts (counts) are linearly correlated with the genes detected by bulk RNA-Seq. (B) The 86 informative genes (SuppTable4) detected by smRNA-FISH show a high dynamic range of transcript expression (counts) across the segmented cells from tissue at W3.5. (C) Frequency density distribution of counts detected in the segmented cells from tissue at W3.5. (D) Range of total transcripts detected in the 22,082 segmented cells. (E) Range of genes (from the 86 used for smRNA-FISH) detected in the segmented cells. In (A), the *x* and *y* axes were $\log_2$-transformed to show the correlation more clearly. In (B) and (C), the transcripts are expressed as $log_{10}(transcripts +1)$ to simplify the visualization of the high dynamic range between genes.

### Supplementary Figure5:

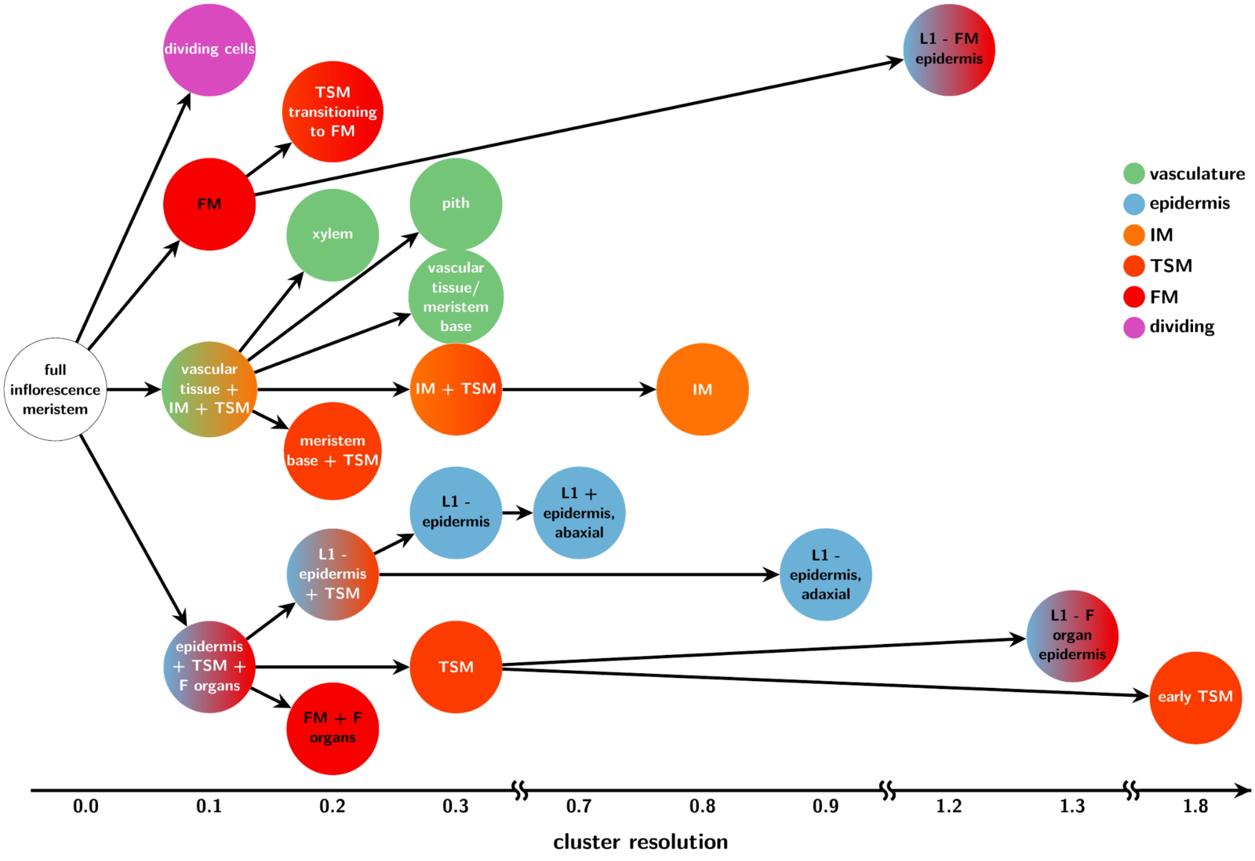

**SupplFig5: Overview of the cell types emerging in the inflorescence meristem depending on the cluster resolution.**

Cell types become distinct depending on clustering resolution parameters. Clusters that separate out at low resolution have very distinct gene expression patterns, whereas those appearing only at high resolutions of the *Seurat* clustering algorithm show less distinct gene expression profiles (IM, inflorescence meristem; TSM, triple spikelet meristem; FM, floret meristem). The color code shows the main cell lineages identified in the plot under specific cluster resolutions.

#### Supplementary Figure6:

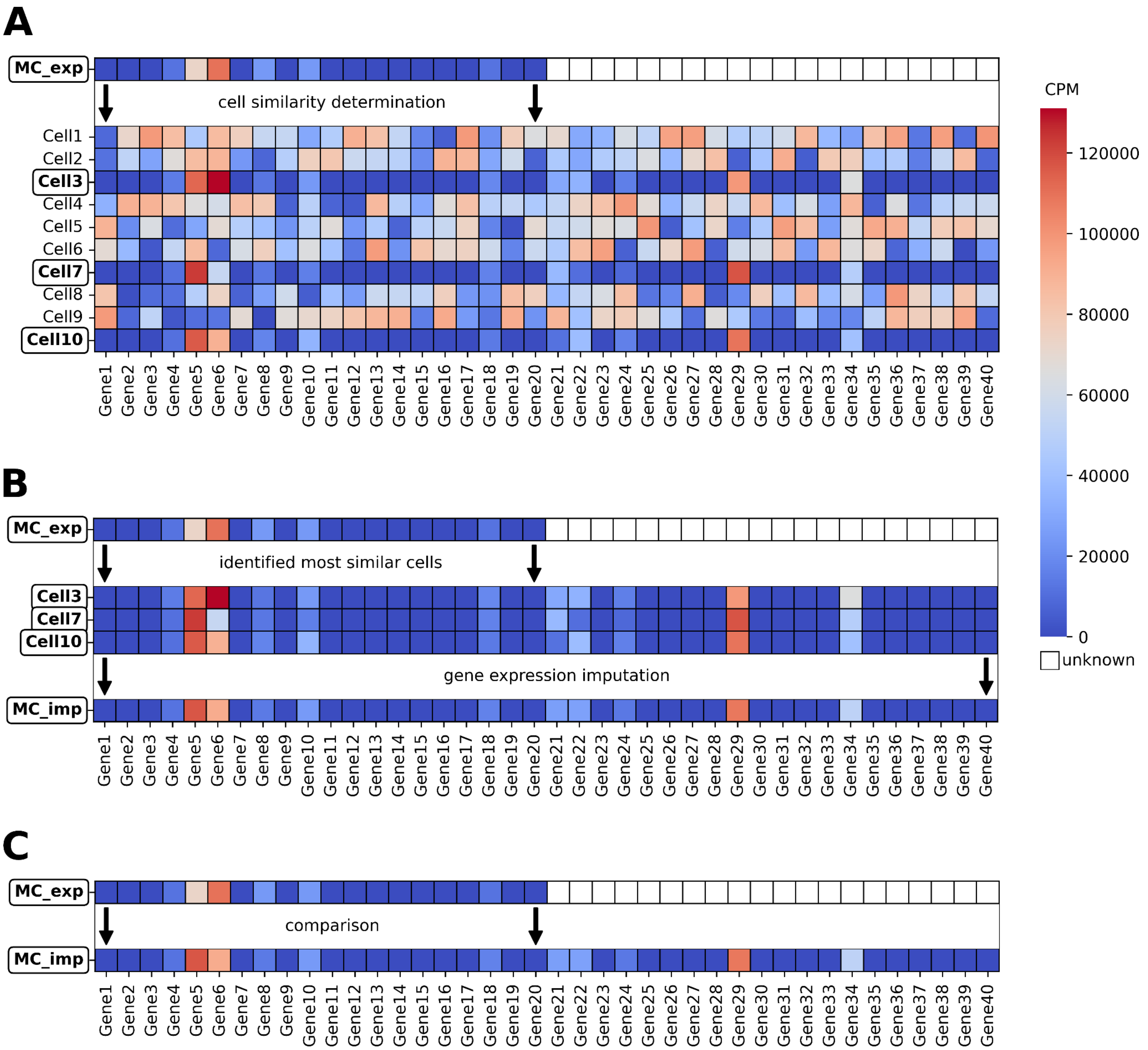

**SupplFig6: Scheme of the gene expression imputation workflow.**

(A) The first step of gene expression imputation is the determination of the most similar scRNA-Seq cells for each cell that was experimentally measured by molecular cartography (MC_exp) based on their gene expression, using the genes from the MC (Gene1-Gene20). Cell similarity is determined by cosine similarity (CS). The most similar cells are highlighted on the left. (B) Gene expression is determined by calculating the weighted average of the gene expression from the most similar cells, using each cell's CS as weight. This yields an imputed version of the same cell from the molecular cartography (MC_imp), containing gene expression information about all genes measured in the scRNA-Seq experiment. (C) Validation by comparing the experimentally determined and computationally imputed version of each MC-cell, using the available genes measured in the MC (Gene1-Gene20). Similarity between imputed and experimentally determined gene expression values was calculated using CS.

79 **Supplementary Figure7:**

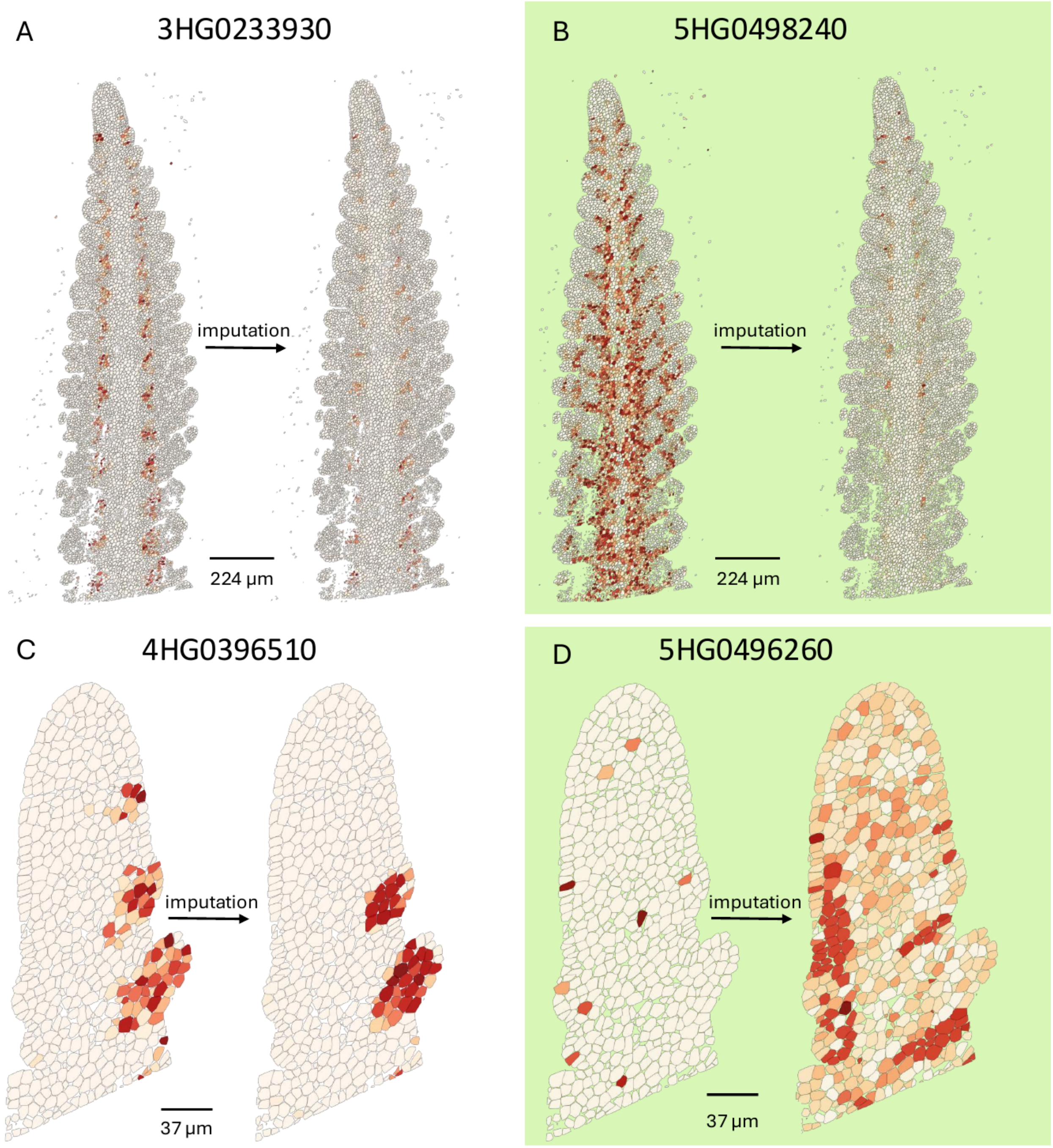


80

**SupplFig7: Comparison of measured and imputed gene expression**

(A, B) The measured and imputed gene expression patterns and levels for two genes in the inflorescence meristem. (C, D) The measured and imputed gene expression patterns and levels for two genes in the vegetative shoot apical meristem. The green background indicates that the genes displayed in (B, D) are affected by the protoplasting procedure, resulting in different patterns.

### Supplementary Figure8:

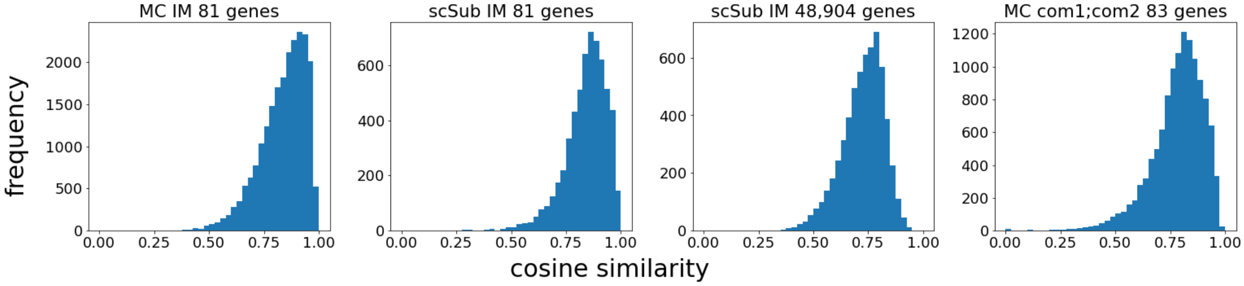

**SupplFig8: Gene expression imputation quality assessment.**

Imputed and experimentally determined gene expression values were compared for four different datasets using cosine similarity (CS) for each cell in the test dataset. The first histogram shows the CS for all 22,082 cells in the inflorescence meristem (IM) Molecular Cartography (MC) dataset for the 81 experimentally determined genes. The second histogram shows the CS for all 5,910 cells in the subset of the scRNA-Seq dataset for the same 81 genes available in the MC. The third histogram shows the CS for the same 5,910 cells in the subset of the scRNA-Seq dataset for all 48,904 genes experimentally determined in the scRNA-Seq experiment. The fourth histogram shows the CS for all 12,092 cells in the *com1a;com2g* MC dataset for the 83 experimentally determined genes.

#### Supplementary Figure9:

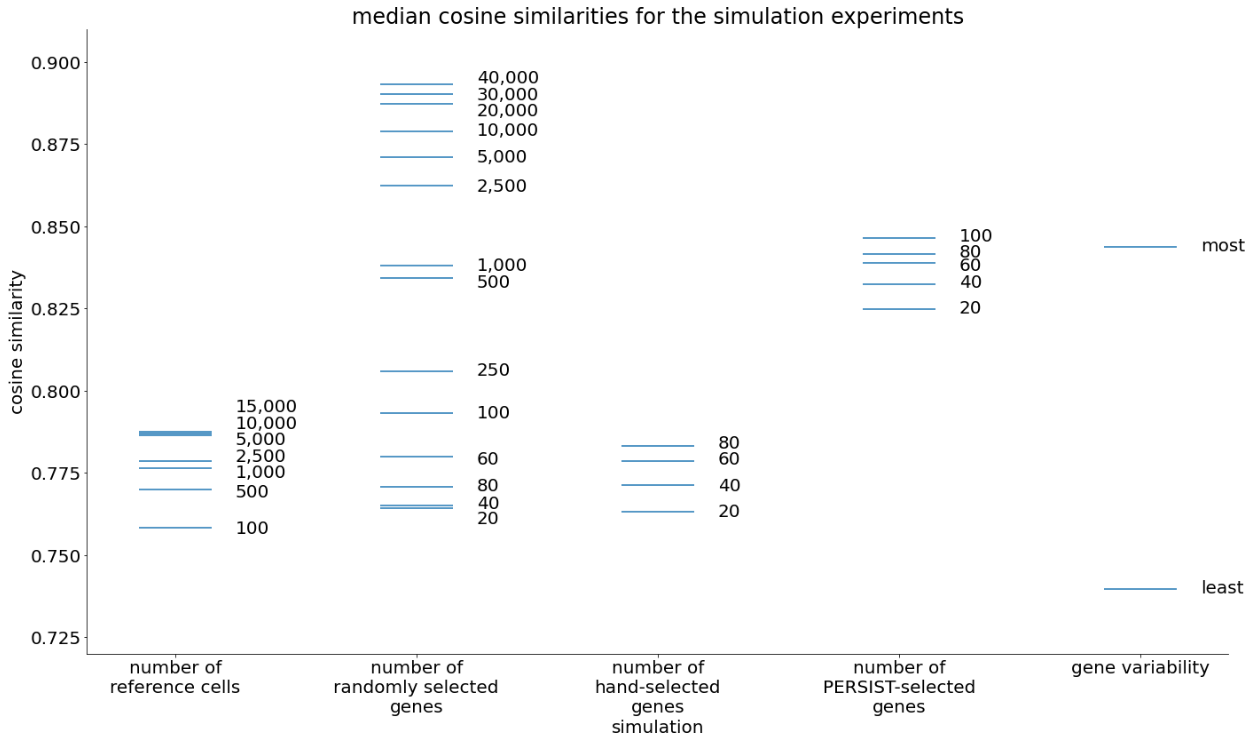

**SupplFig9: Gene expression imputation quality simulation.**

Imputation quality for five different simulations with different numbers of reference cells, for different numbers of genes selected for the similarity determination, either by random selection, hand selection of known marker genes and genes of interest, by selection with the *PERSIST* tool, or by selecting the top and bottom 250 most and least variable genes. Each line represents the median cosine similarity score from the validation, which compares imputed gene expression patterns to the experimentally measured patterns. The displayed medians each represent a mean value from n = 5 simulation experiments.

114 **Supplementary Figure10:**

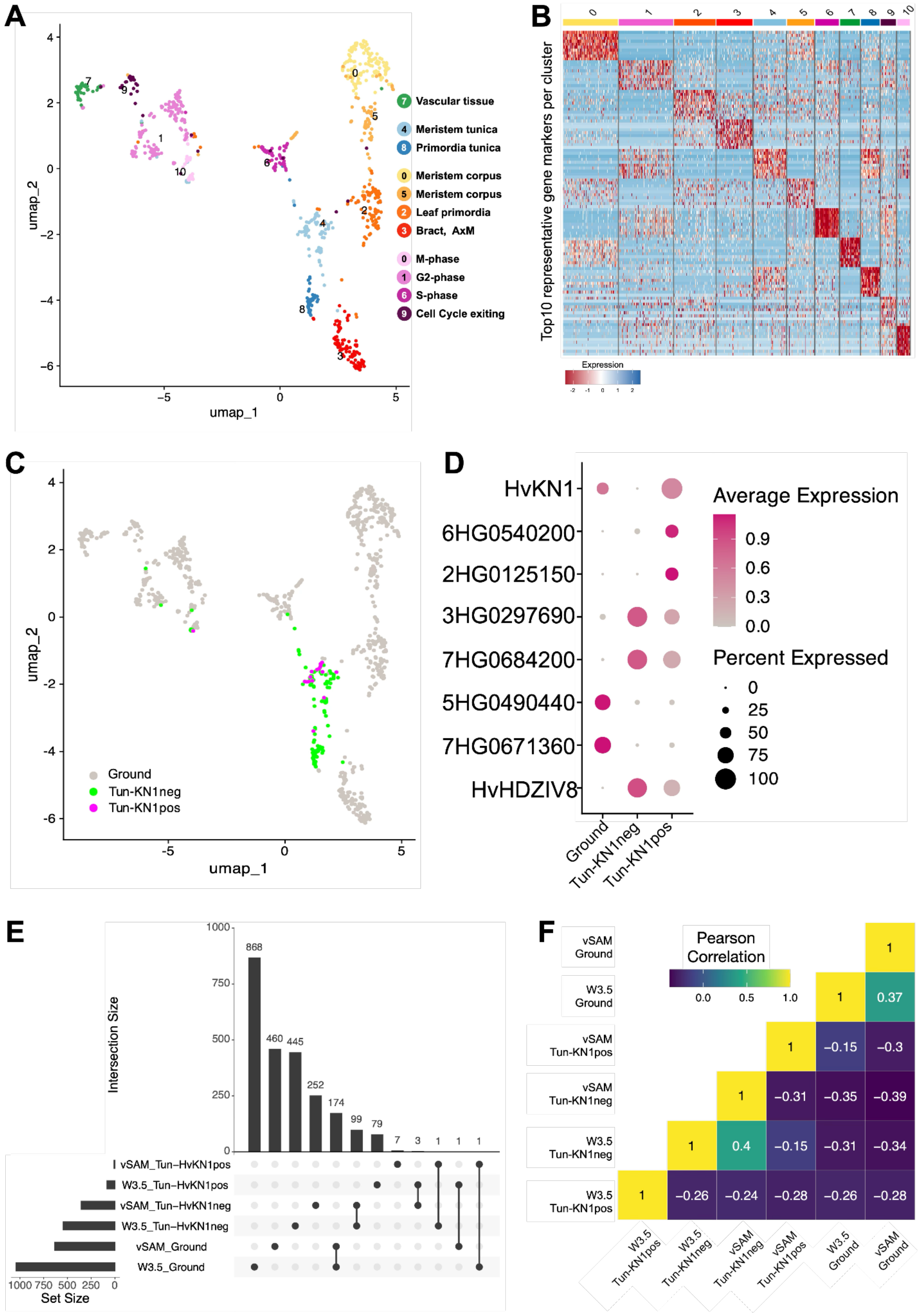


115

**SupplFig10: Single-cell atlas of barley vegetative SAM.**

(A) UMAP dimensional reduction of scRNA-Seq in the vegetative SAM (vSAM). (B) Top 10 marker genes per cluster prioritized by $P_{adj}$. (C) *HvKN1* expression in tunica cells. (D) Selected marker genes in tunica cells expressing *HvKN1* in the vSAM. (E) UpSet plot showing the number of differentially expressed genes (DEGs) specifically identified or shared between cell populations: tunica expressing *HvKN1*, tunica lacking *HvKN1* expression, and ground tissue (corpus), each for vSAM and W3.5. Set size indicates the total number of DEGs identified in each group of cells. Intersection size shows the number of DEGs shared between the indicated cell groups. (F) Pearson correlation between the number of DEGs identified and the number of shared DEGs comparing all different cell populations. Scale bars: 50 µm.

### Supplementary Figure11:

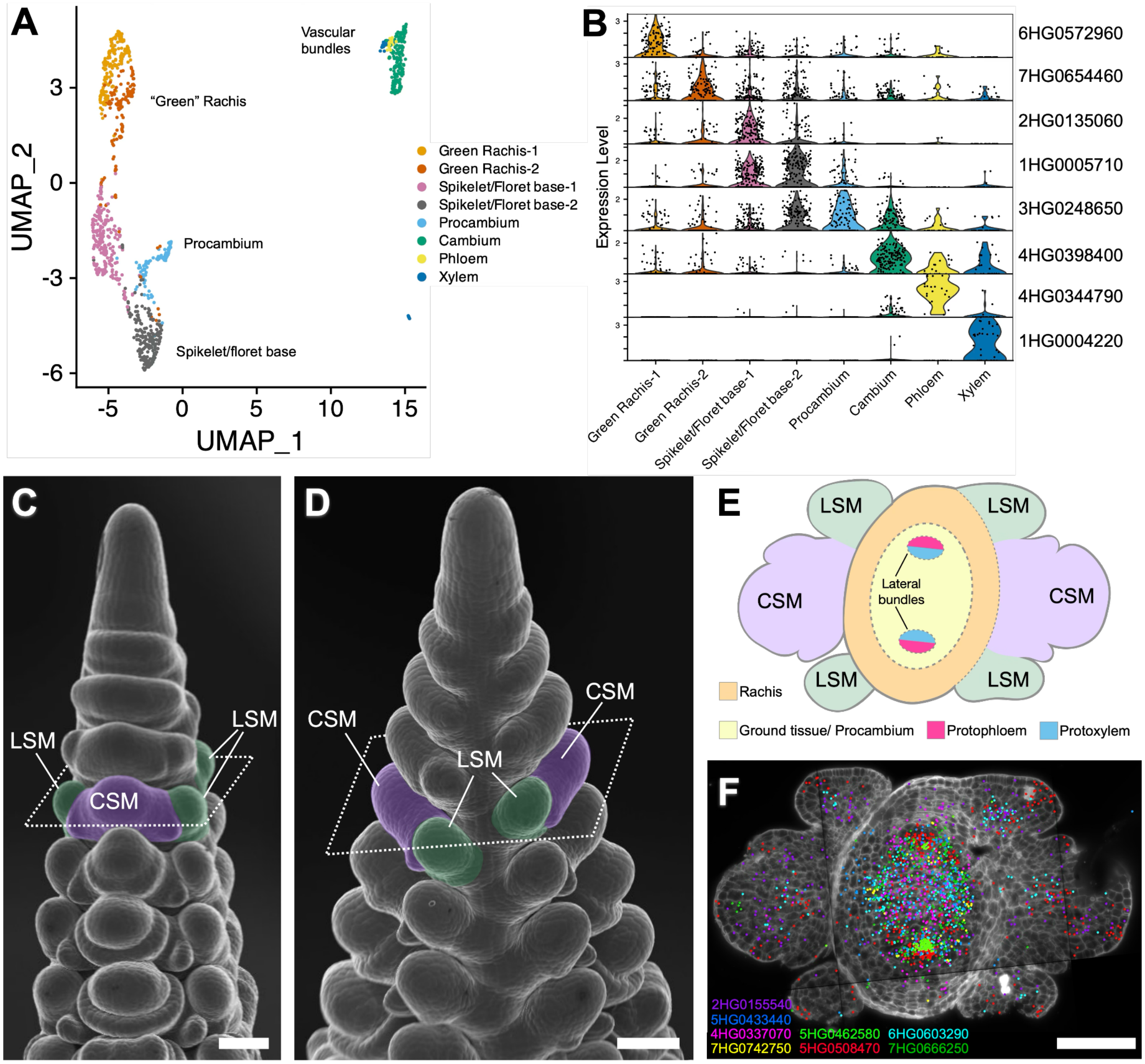

**SupplFig11: Subcluster analysis in vasculature.**

(A) UMAP plot with subcluster organization in vascular cells. (B) Violin plots of selected markers expressed in subclusters. (C) Scanning electron micrograph frontal view of a spike, indicating the position of the transverse section (dashed lines) used in multiplex smRNA-FISH (E,F) for vascular cells (LSM in green, CSM in violet). (D) Lateral view of the same spike. (E) Schematic representation of transversal section at the position indicated in (C) and (D). (F) Selected expression profiles in vascular tissues based on multiplex smRNA-FISH. Scale bars: 100 μm.

**Supplementary Data:**

**Identification of cluster and marker genes for spike greening**

The total number of grains produced on a spike depends on the number of SMs that are initiated, and on their maintenance. SM number is largely controlled by flowering time genes that set heading time, whereas the development of the vasculature, light signals and activation of chloroplasts promotes SM survival {Huang, 2023 #1530}. In developing spikes of the subfamily Pooideae, development of plastids and the photosynthetic machinery in the rachis can be observed as an acropetal gradient of green cells in the vasculature. Using the identified clusters related to vascular tissues, we analysed these cells by unbiased subclustering (**SupplFig11, SupplFig12**). We identified marker genes such as *Thioredoxin z* (*HvTRX*, 4HG0341820) involved in plastid division, and *PHOTOSYSTEM II LIGHT HARVESTING COMPLEX GENE 2* (*HvLHCB2*;*3*, 5HG0499400) and Photosystem I reaction center subunit N (*HvPSAN*, 2HG0113750), both involved in photosynthesis and important for green rachis pigmentation.

At W3.5, the developing spike forms lateral bundles in the rachis, which will connect the vascular tissues to the spikelets. In our subclusters, we can recognize the procambium in the ground tissues and the cambium differentiating into protoxylem and protophloem, patterning the lateral bundles. We identified additional protophloem and protoxylem markers. Among the new markers for developing barley phloem, we found *HvBAM5* (4HG0344790), DNA-BINDING WITH ONE FINGER (DOF) transcription factor genes (5HG0441840, 5HG0526260) and two homologs of Arabidopsis *ALTERED PHLOEM DEVELOPMENT* (6HG0570890, 7HG0738570), and for the developing xylem we found the CLE-peptide homolog of FON2 SPARE1 in barley (*HvFOS1*, 1HG0008860), six laccases (7HG0667290, 4HG0357240, 3HG0302570, 1HG0073500, 2HG0135650, 3HG0302520), two LOGs (4HG0408000, 1HG0043530), and two basic helix-loop-helix transcription factors (7HG0731430, 5HG0484930) related to *PERICYCLE FACTOR TYPE-A 2* (*PFA2*) and *TARGET OF MONOPTEROS 5*, respectively, already associated with xylem differentiation in *Pinus* and Arabidopsis (**SupplTable13**).

#### Supplementary Figure12:

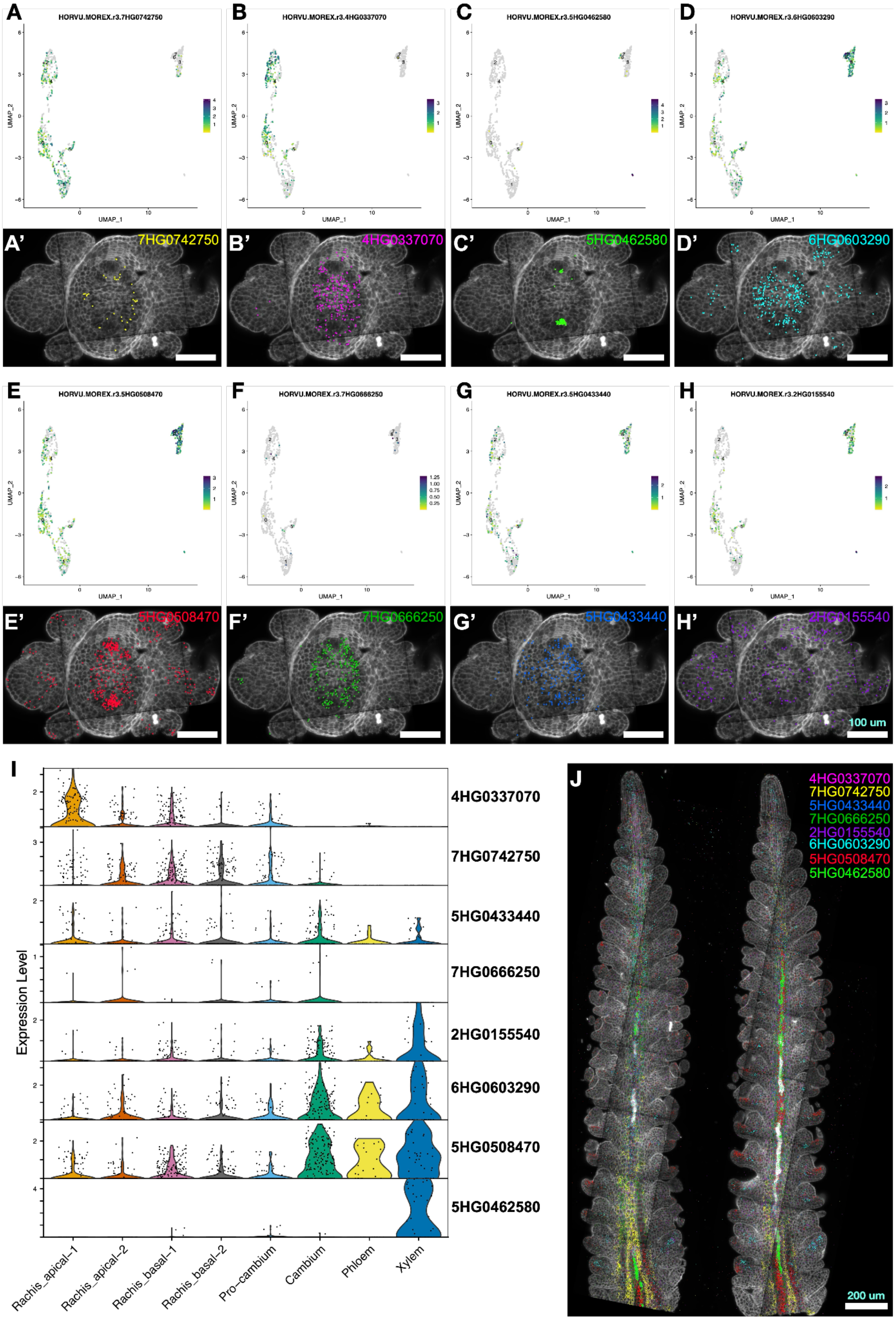

**SupplFig12: Subcluster analysis and validation in vascular tissues**

(A-H) UMAPs in subcluster analysis using clusters related to vascular tissue and their smRNA-FISH expression profiles in transverse sections of W3.5 developing spikes in barley. (I) Violin plot of gene expression in subclusters. (J) Expression profiles in sagittal sections of developing spikes. Scale bars: (A′)–(H′) 100 μm; (J) 200 μm.

### Supplementary Figure13:

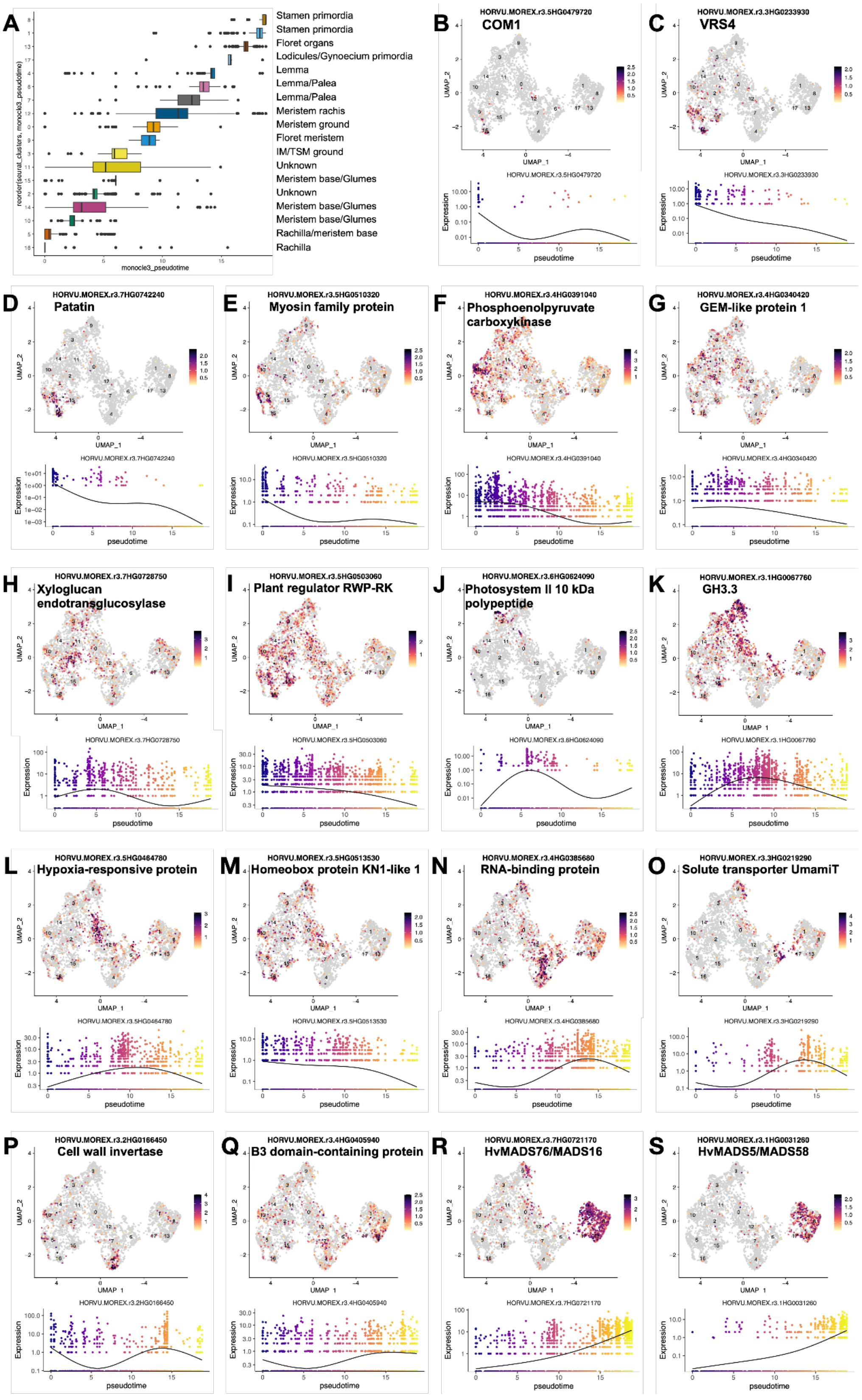

**SupplFig13: Subcluster analysis in corpus cells and their modulation across the pseudotime.**

(A) Corpus subclusters arranged along pseudotime. (B) Expression levels of *COM1* in corpus subclusters and its modulation in pseudotime. (C) Expression levels of *VRS4* in subclusters and its modulation in pseudotime. *COM1* and *VRS4* are early markers for spikelet meristem formation. (D)-(S) Expression profiles and their modulation along pseudotime for selected marker genes in corpus subclusters.

### Supplementary Figure14:

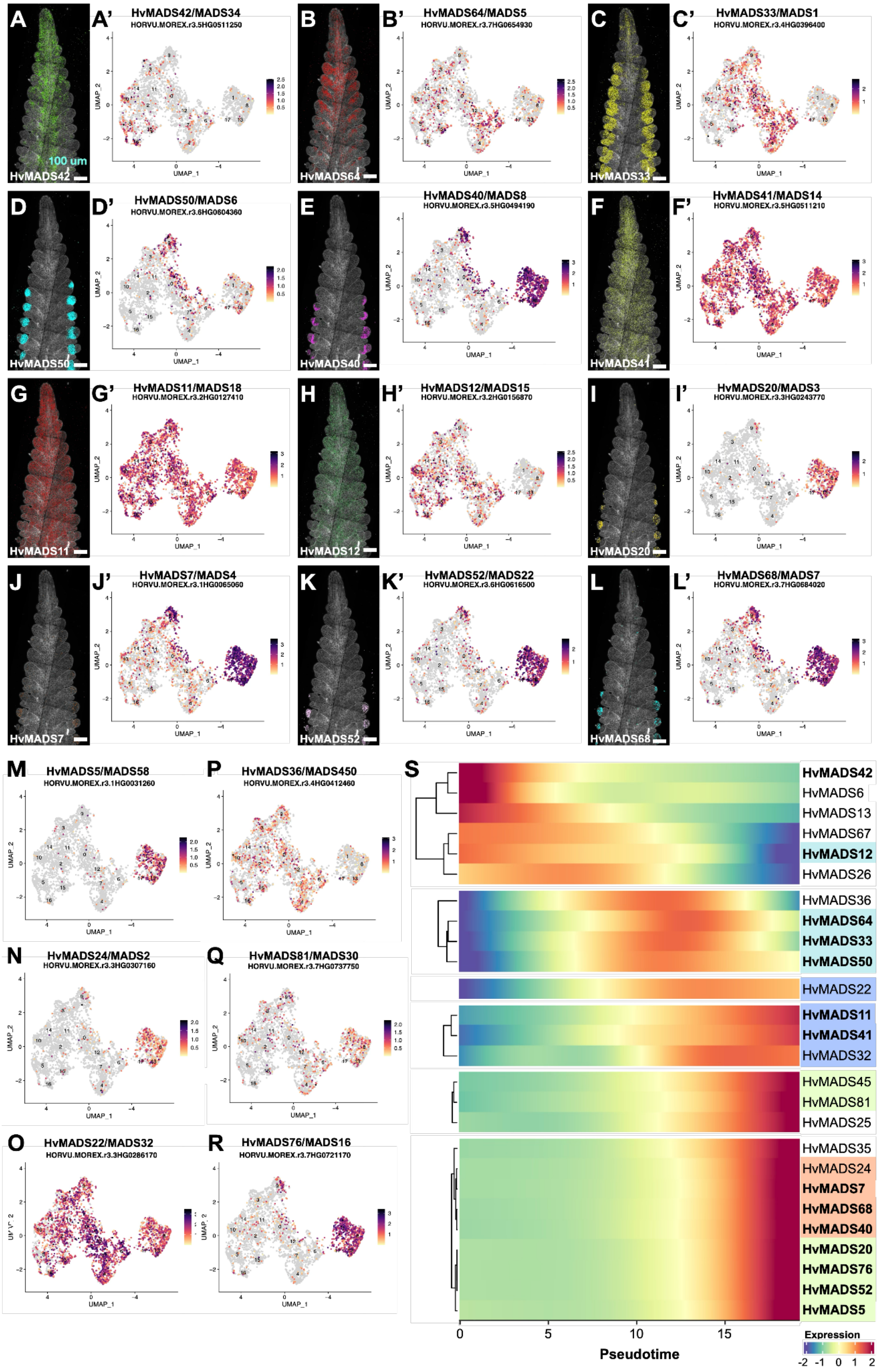

**SupplFig14: MADS-box transcription factor genes expressed in corpus subclusters and their modulation along pseudotime**

(A)–(E) MADS-box smRNA-FISH expression patterns and their expression profiles in corpus subclusters. (F)–(L) Additional expression patterns of selected MADS-box genes. (M)–(R) Additional UMAP plots of MADS-box genes expressed in corpus subclusters. (S) Modulation along pseudotime of MADS-box transcription factor genes expressed in corpus subclusters. MADS-box genes with smRNA-FISH data reported in this study are shown in bold. Assignment of MADS-box classes: A, lilac; B, peach; C, pale green; E, pale blue. Scale bars: 100 μm.

#### Supplementary Figure15:

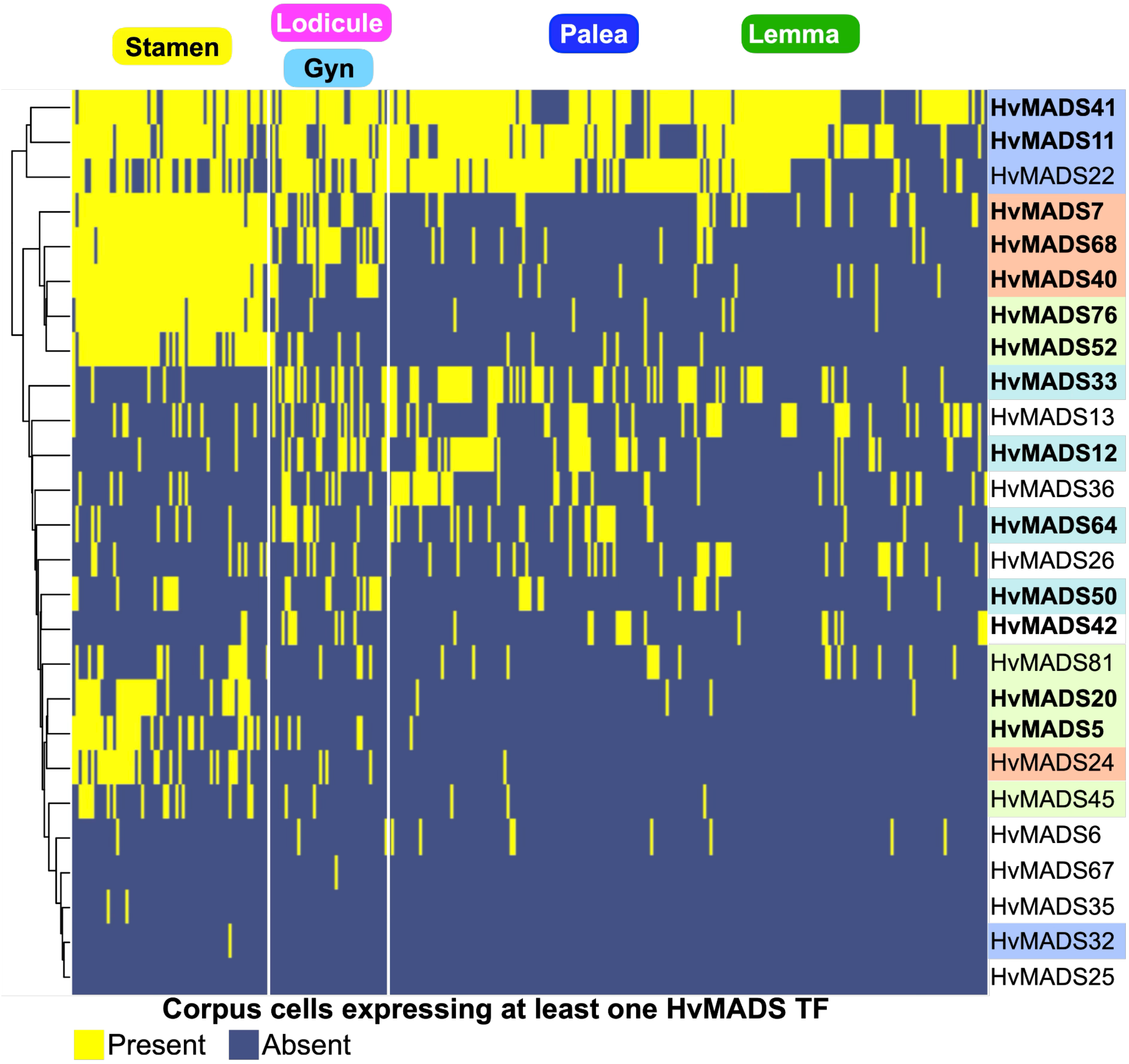


**SupplFig15: MADS-box transcription factor genes co-expressed in corpus cells.** The presence or absence of MADS-box transcription factor genes detected by scRNA-Seq in corpus cells, hierarchically ordered by the MADS-box gene expressed in each cell. Assignment of MADS-box classes: A, lilac; B, peach; C, pale green; E, pale blue. The origin of the putative floret organ of the cells is indicated above the heat map, according to the HvMADS class.

**Supplementary Figure16:**

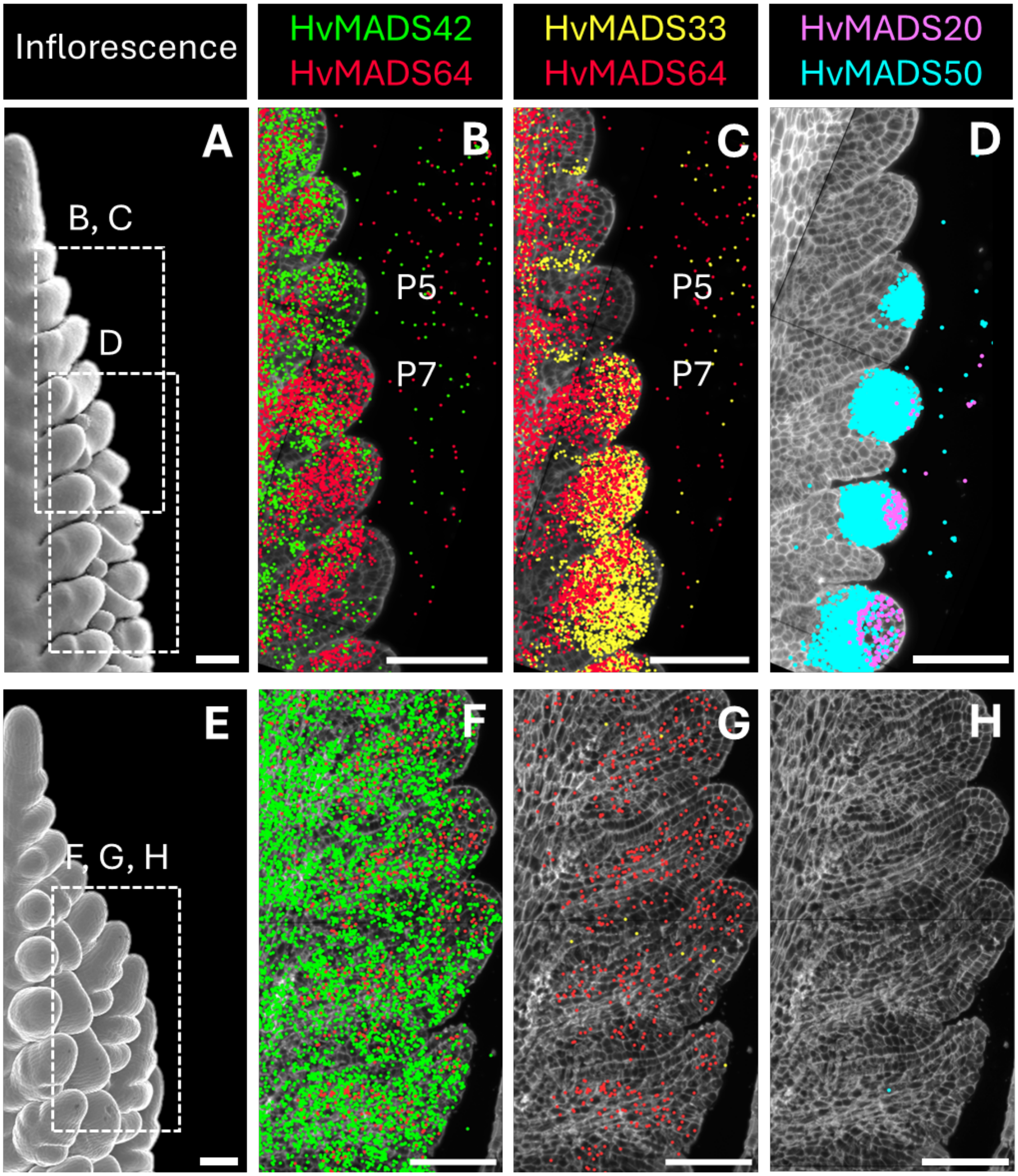

**SupplFig16: Stage-specific MADS-box transcription factor genes expressed in WT and *com1a;com2g* inflorescences.**

(A) Scanning electron micrograph of WT barley inflorescence at W3.5. Dashed areas indicate the location of primordia displayed in panels B–D. (B,C) Longitudinal sections of developing WT spikelets: smRNA-FISH data showing the localisation of *HvMADS42* (green), *HvMADS64* (red), and *HvMADS33* (yellow) transcripts. (D) Developing WT florets: smRNA-FISH data showing the localization of *HvMADS20* (purple) and *HvMADS50* (cyan) transcripts. (E) Scanning electron micrograph of *com1a;com2g* branched inflorescence. The segmented square indicates the location of IM-like branches displayed in panels F-H. (F-H) Longitudinal sections of *com1a;com2g* IM-like branches: smRNA-FISH data showing the localization of *HvMADS42* (green), *HvMADS64* (red), *HvMADS33* (yellow), *HvMADS20* (purple), and *HvMADS50* (cyan) transcripts. Scale bars: 100 µm.

#### Supplementary Tables

**Supplementary Table 1.** Metrics summary for the scRNA-Seq libraries from the different replicates used in this study.

**Supplementary Table 2.** Differentially expressed genes during the protoplasting process ($-3 \le log_2FC \ge 3$, $FDR < 1x10^{-5}$).

**Supplementary Table 3.** Unique markers per cluster in developing spikes at WS3.5 (pct1 > 0.25; pct2 < 0.3; $p_{adj} < 0.001$).

**Supplementary Table 4**. Reported marker genes used to annotate cluster/subcluster identities and their closest homologs in maize and Arabidopsis.

**Supplementary Table 5**. Genes used for multiplex fluorescence in situ hybridization.

**Supplementary Table 6.** Annotation of MADS-box transcription factor genes in barley and their putative orthologs in rice by reciprocal BLAST.

**Supplementary Table 7.** Unique markers per cluster in the vegetative SAM (pct1 > 0.25; pct2 < 0.3; $p_{adj} < 0.001$).

**Supplementary Table 8.** Statistical values describing the gene expression imputation results.

**Supplementary Table 9.** Gene markers for ground tissue and tunica cells positive or negative for *HvKN1* expression in developing spikes at WS3.5.

**Supplementary Table 10.** Gene markers for ground tissue and tunica cells positive or negative for *HvKN1* expression in the vSAM.

**Supplementary Table 11.** Gene markers for TSM founder and primordia, and the rest of corpus cells (ground). $P_{adj} \le 0.05$.

**Supplementary Table 12.** Imputed differentially expressed genes between IM, founder and primordial cells using MAST ($p_{adj} < 0.05$, pct.1/pct.2 ratio ≥ 1.2).

**Supplementary Table 13.** Unique markers per subcluster in vascular tissues (pct1 > 0.25; pct2 < 0.3; $p_{adj} < 0.001$).

**Supplementary Table 14.** Unique markers per subcluster in corpus tissues (pct1 > 0.25; pct2 < 0.3; $p_{adj} < 0.001$).

**Supplementary Table 15.** MADS-box transcription factor transcripts detected by scRNA-Seq.

**Supplementary Table 16.** Modulated genes in pseudotime analysis (Morans I > 0.1; q ≤ 0.01).

**Supplementary Table 17.** Imputed differentially expressed genes along spikelet development (P1–P19).

**Supplementary Table 18.** Imputed differentially expressed transcription factor genes marking spikelet development (P1–P19).

**Supplementary Table 19.** Imputed differentially expressed genes in cells of regions in the floret meristem central zone (FMc), abaxial (FMab) and adaxial (FMad) using MAST ($p_{adj} < 0.01$, pct.1/pct.2 ratio > 1).

**Supplementary Table 20.** Genes selected for the P1–P19 barcode and the WT *com1a;com2*g barcode.

**Supplementary Table 21.** Imputed differentially expressed genes between IM and FM in WT and between IM and the equivalent plastochron by position in *com1a;com2g*.

**Supplementary Table 22.** Description of anchors found during the integration of the inflorescence meristem and the vegetative shoot apical meristem, as well as imputation quality results.
